# Mitophagy modulation rescues single large-scale mitochondrial DNA deletion (SLSMD) disease symptoms in the *C. elegans uaDf5* animal model

**DOI:** 10.1101/2024.10.27.620333

**Authors:** Ryan M. Mendel, Sofie Matsuno, Alex Lu, Marni J. Falk, Suraiya Haroon

**Author notes:** Corresponding Author: Children’s Hospital of Philadelphia Abramson Pediatric Research Center 3615 Civic Center Blvd, Suite 1002E Philadelphia, PA 19104.

## Abstract

<u>S</u>ingle large <u>s</u>cale <u>m</u>itochondrial DNA (mtDNA) <u>d</u>eletions (SLSMD) underlie a range of sporadic or maternally inherited primary mitochondrial diseases having significant morbidity and mortality, including Pearson syndrome, Kearns-Sayre Syndrome, or Chronic Progressive External Ophthalmoplegia. Therapeutic development has been hindered by limited existing knowledge on mtDNA quality control and a lack of SLSMD animal models. To address this challenge, we utilized the *C. elegans* heteroplasmic SLSMD strain, *uaDf5,* to objectively screen for potential therapies. As mitophagy modulation has been implicated in mtDNA homeostasis, we screened a library of mitophagy modulating compounds to determine their comparative effects to rescue mitochondrial unfolded protein (UPR^mt^) stress induction in in *uaDf5* SLSMD worms. Interestingly, Thiamine was discovered to be an effective positive control, significantly reducing mitochondrial stress in this model. Two lead therapeutic candidates from the mitophagy library screen were Hemin and Celastrol (Tripterin). Celastrol is a mitophagy activating anti-inflammatory and metabolic modifying natural product derived compound, that rescued multiple fitness outcomes (thrashing, development, survival) and reduced the mitochondrial stress in *uaDf5* animals in a mitophagy-dependent fashion. This study highlights the utility of the *uaDf5* worm model to enable preclinical identification of therapeutic candidate leads for SLSMD-based heteroplasmic mtDNA diseases and identifies possible therapeutic candidates that serve as mitophagy modulators to improve health and specifically reduce heteroplasmy levels in SLSMD diseases.

## INTRODUCTION

Mitochondria are specialized energy producing organelles that harbor their own mitochondrial DNA (mtDNA) genome. The mtDNA codes for 13 proteins essential for bioenergetics and as such, maintaining mtDNA integrity is paramount to organismal fitness^1,2^. However, mtDNA mutations can arise sporadically with aging, during replication, or exposure to toxins, which can give rise to primary mitochondrial diseases (PMD)^2,3^. One class of PMDs that arises from a specific subtype of mtDNA mutations the either arises *de novo* or inherited through the maternal lineage are <u>s</u>ingle large <u>s</u>cale <u>m</u>tDNA <u>d</u>eletions (SLSMD) syndromes^2,4–6^, which include Kearns-Sayre Syndrome (KSS)^7^, Pearson Syndrome (PS)^8^, and Chronic Progressive External Ophthalmoplegia (CPEO)^9^. As a multicopy genome (10s to 1000s per cell), SLSMD and wild-type mtDNA genotypes co-exist within the same cell, a state defined as heteroplasmy. In instances where the levels of mutated genomes can no longer be compensated for by the wild-type copies, the clinical threshold of reduced mitochondrial respiratory capacity is crossed and disease symptoms manifest. The severity of these syndromes often correlates to the level of SLSMD heteroplasmy in specific tissues, as well as the size of the deletion^4^. SLSMD diseases collectively represent a class of rare disease that currently remains incurable, with high morbidity and mortality across the lifespan.

Developing therapeutics for SLSMD disorders is hindered by limited understanding both of DNA repair mechanisms within the mitochondrial matrix that respond to loss mtDNA fidelity, as well as the ways in which wild-type mtDNA genomes are distinguished from mtDNA genomes harboring large-scale deletions^10,11^. One cellular mechanism that has been shown to modulate mtDNA heteroplasmy is mitophagy^12–14^, whereby dysfunctional mitochondria are targeted for degradation using the autophagy machinery. Therefore, a plausible therapeutic opportunity in SLSMD disorders may be to modulate mitophagy to reduce the mutant mtDNA genome heteroplasmy load. In fact, the regulation of mitophagy has been shown to be a potential therapy for both primary and secondary forms of muscle-based mitochondrial dysfunction^15–17^. Despite these promising results, SLSMDS, and PMDs in general, currently have no cure or FDA-approved therapies due to limitations in available biological models in which to test empiric candidates or screen compound libraries. In large part, this stems from the technology to develop targeted mtDNA deletions being in its infancy and, of particular importance, biologic limitations whereby transmittance of the mutant mtDNA genome to the next generation has been found to be unstable^18,19^.

A naturally occurring *C. elegans* mutant strain, *uaDf5,* has been identified to harbor a 3.1 kb heteroplasmic large-scale mtDNA deletion that encompasses 7 tRNA and 4 mRNA genes^20–27^. In this study, several substrains of *uaDf5* with varying heteroplasmy levels were established and characterized for both gross animal phenotypes and effects on mitochondrial physiology. These strains exhibited relevant subcellular and organismal defects akin to problems reported in SLSMD patients. Exploiting their increased mitochondrial stress phenotype, Thiamine was identified as an effective positive control. Subsequently, a targeted screen of 62 mitophagy modulating drugs was performed in *uaDf5,* which identified 2 mitophagy activators as promising therapeutic hits. Developing this SLSMD model enables larger-scale screening of potential therapeutic compounds for mtDNA diseases, which can be validated in human cell samples to streamline development of novel therapeutic leads that may ultimately improve the health outcomes and survival of SLSMD patients.

## METHODS

### Strains

All strains were maintained at 20°C and grown on nematode growth media (NGM) plates seeded with OP50 *E. coli*^28^. The original N2 (wildtype), *hsp-6p::GFP* (SJ4100)^29^, *myo-2p::mCherry* (VS21 - deposited by Ho Yi Mak), *cox-4(zu476[cox-4::eGFP::3xFLAG])* (JJ2586 – deposited by Michael Eastwood) and *uaDf5;him-8(e1289)* (LB138)^20^ strains were obtained from the Caenorhabditis Genetics Center (CGC) at the University of Minnesota. The *him-8(e1489)* mutation was crossed out of the LB138 strain to obtain a strain where the only mutation in the *uaDf5* strain is the mtDNA deletion. The *uaDf5*; *hsp-6p::GFP* #1 and #2 strains were derived by mating the *uaDf5* hermaphrodites and males generated from the SJ4100 strain. The *uaDf5*; *myo-2p::mCherry; hsp-6p::GFP* (Low and Medium Δ %) strains were derived using *uaDf5* hermaphrodites and males from the cross of the SJ4100 and VS21 strain. The *uaDf5*; *myo-2p::mCherry; hsp-6p::GFP* High Δ % arose spontaneously with higher heteroplasmy from *uaDf5*; *myo-2p::mCherry; hsp-6p::GFP* Medium Δ % strain. Worms were grown in 20°C on NGM (Nematode Growth Media) with OP50 bacteria per standard practice^30^.

### mtDNA Quantification and Sequencing

#### DNA extraction

For single worm or population heteroplasmy assessment, single adult worm or 30 adult worms were places into individual tubes of a PCR strip tube with 8 µl or 15 µl of extraction reagent (Quantabio, 97065-350). The samples were incubated at 95°C for 30 minutes, and then, 8 µl or 15 µl of stabilization buffer (Quantabio, 97065-350) was added to neutralize the reaction. The samples were used immediately since storage at either −20°C or −80°C led to mtDNA degradation as assessed by qPCR.

#### qPCR

Each sample was tested with three primer sets. The first primer set (cel-ND5) amplifies mtDNA sequence away from the *uaDf5* deletion to quantify total mtDNA count. The second primer set from Liau *et al.* hybridize with sequence flanking the *uaDf5* deletion, which amplifies sequence across the deletion to quantify mutant mtDNA. The third primer set (*ges-1*) amplifies nuclear sequence to normalize data acquired for mtDNA. Per sample, 3 different reactions using the 3 different primer sets were set up. Per reaction, 50µl reaction was prepared with 5 µl of DNA, 25 µl of Powerup Sybrgreen Mastermix (Applied Biosystems, A25742), 1µl of forward primer, 1 µl reverse primer, and 18 µl of molecular grade water (ThermoFisherScientific, BP281910). Each reaction was split into 3 wells of 15 µl to attain three technical replicate data per sample. The 384-well plate (Applied Biosystems, 4309849) was sealed with microamp optical adhesive film (Applied Biosystems, 4311971). The samples were run in the ViiA 7 Real-Time PCR System (Applied Biosystems) with the cycling conditions of 1x (2 min at 50°C), 1x (2 min at 95°C), and 40x (3 sec at 95°C and 30 sec at 60°C).

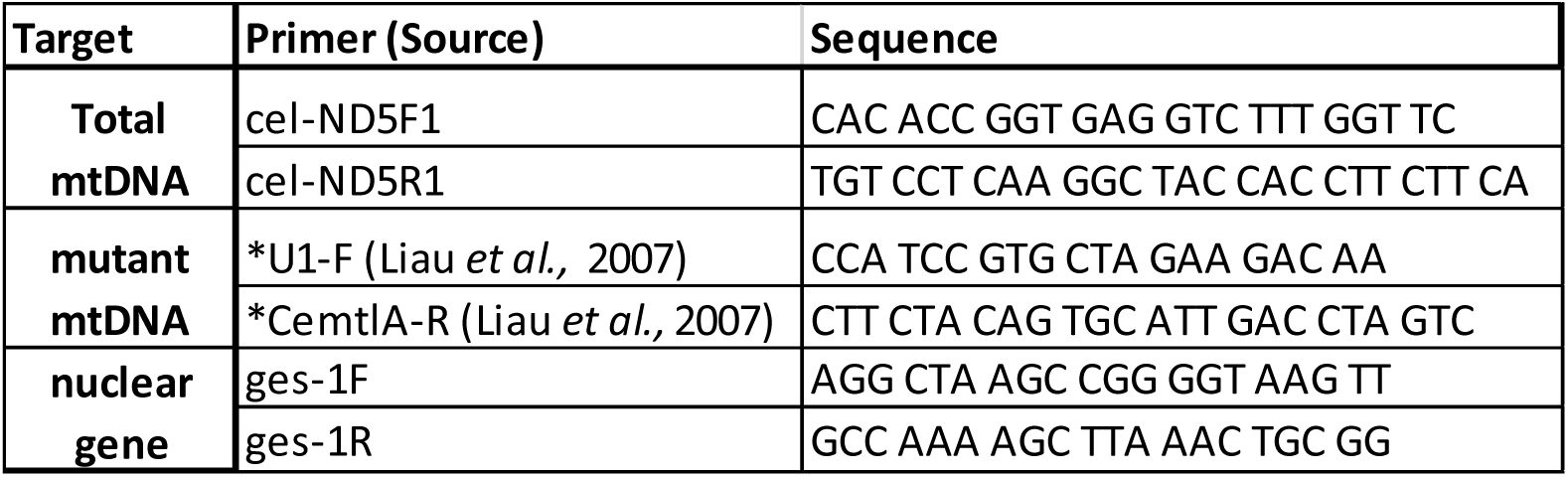

The cycle threshold (CT) values of the 3 technical replicates were averaged and then the total and the mutant mtDNA levels were calculated using the formula 2^-(CT_mtDNA_-CT_nDNA_). The heteroplasmy of the mutant mtDNA was assessed using the formula 100*CT_mutant_/CT_total._ At least three separate biological replicate samples were collected per sample. Student t-test analysis was performed to determine statistical significance in GraphPad Prism version 10.0.0 for Windows, GraphPad Software, Boston, Massachusetts USA, www.graphpad.com.

#### Sequencing

10 µl of a qPCR sample with *uaDf5* DNA source with the mutant mtDNA primer set^27^ was inactivated by adding 4 µl of ExoSAP-IT PCR Product Cleanup Reagent (ThermoFisherScientific, 78201.1.ML) and incubating at 37°C for 15 min and 80°C for 15 min. The inactivated sample combined with either the forward or reverse primer was sent to Genewiz (Azenta Life Sciences, NJ) for sequencing. Sequence alignment analysis was performed using ApE^31^.

### Developmental Assay and Hatch Rate

To assess development, 50 embryos were picked onto a 3.5cm petri dish, and after 4 days, the hatched worms were counted and assessed for life stage. In the case of drug treatment, 150-200 µl of solution containing either DMSO only or Celastrol in DMSO was applied to the plate, spread, and dried prior to adding the embryos. At least 3 biological replicates were completed per condition with 50 embryos being picked for each replicate. Student t-test analysis was performed to determine statistical significance in GraphPad Prism version 10.0.0 for Windows, GraphPad Software, Boston, Massachusetts USA, www.graphpad.com.

### Synchronizing Worms via Bleaching

Worms were washed off from plates and transferred into 15 ml conical tubes using 4 ml S basal medium (23.4 g NaCl, 4 g K_2_HPO_4_, 25 g KH_2_PO_4_, 4 ml cholesterol (5 mg/ml in ethanol), 4 L milliQ water, pH = 7). 1 ml of 100% fresh bleach (Clorox, 32263) and 500ul 5 M NaOH was added to the worms. The tubes were gently agitated for 2-5 min, until all worm bodies disintegrated. The tubes were then centrifuged for 1 min at 182 rcf in the Eppendorf centrifuge 5810 R, the supernatant was removed, the pellet was washed with 10 ml of fresh S. basal and centrifuged again. The supernatant was removed, the pellet was washed with 10 ml S basal medium and centrifuged again. Most of the supernatant was removed and the pellet of embryos were resuspended by vortexing in the remaining S basal medium to be placed onto plates for experiments.

### Fecundity Assay

On day 0, single L4 parent worms were placed on a fresh plate and the worm was transferred onto a fresh plate every day for 3 more consecutive days and on day 4, the parent worm was killed. Hatched progeny was counted two days after the adults were removed and the progeny was summed to assess the fecundity per worm. At least three biological replicates were completed with a minimum of 5 worms assessed per replicate. Student t-test analysis was performed to determine statistical significance in GraphPad Prism version 10.0.0 for Windows, GraphPad Software, Boston, Massachusetts USA, www.graphpad.com (Dotmatics). In the case of drug treatment, the plates were prepared by adding 150-200 µl of solution containing either DMSO or Celastrol in DMSO to the plate, spread, and dried. Then embryos were added to the plate and grown for 3 days to obtain treated L4 animals, at which point, the assay was conducted as described. Student t-test analysis was performed to determine statistical significance in GraphPad Prism version 10.0.0 for Windows, GraphPad Software, Boston, Massachusetts USA, www.graphpad.com.

### Thrashing Assay

L4 animals, or day 0 of adulthood, were collected and aged 5 or 7 days, at which point 24 animals were picked into 24 individual wells in 100 µl of S basal medium per well of a 96-well clear u-bottom well plate (Evergreen Labware, 222-8032-01R). Then the animals were imaged for 6 seconds at 16.7 frames per second using the Basic Imaging Platform developed by Tau Scientific Instruments LLC. The recording was analyzed to measure body bends per second (bbps) using the open-source software ImageJ^32–34^ with the plugin “wrmtrck”^35^. BBPS per worm was plotted for each genotype. Each biological replicate had 24 animals and at least 3 biological replicates were performed per condition. Statistical analysis was done using the student’s t test analysis in GraphPad Prism version 10.0.0 for Windows, GraphPad Software, Boston, Massachusetts USA, www.graphpad.com.

### RNA Interference (RNAi)

Agar plates for RNAi experiments were made following the standard NGM protocol^30^ with the addition of 25 mg/ml carbenicillin and 1 M IPTG in the agar plate spread with RNAi bacteria. RNAi bacterial cultures were grown in LB with 5 mg/ml tetracycline and 25 mg/ml carbenicillin overnight in at 37°C and concentrated 10-fold prior to seeding onto agar plates. Once the plates were dry, embryos collected via bleaching were placed on the RNAi plates and grown for 3 or 4 days prior to mitochondrial stress and mtDNA quantification.

### Biosorter Assessment of Mitochondrial Stress and Worm Length

Mitochondrial stress was assessed using the *hsp-6p::GFP* construct, where the *hsp-6* promoter, activated upon mitochondrial unfolded protein stress (UPR^mt^), drives GFP expression thereby allowing assessment of mitochondrial stress via quantification of fluorescence. Large quantities of synchronized animals collected via bleaching was grown on conditions of choice for 3-4 days. Then the worms were washed off the plates using S. Basal and collected into 50 mL conical tubes. The fluorescence was measured on the BioSorter® Large Particle Flow Cytometer from Union Biometrica using the FlowPilot™ software. First, any object outside the normal size range for L4 to adult worms were censored. To evaluate the level of UPR^mt^, the green fluorescence was normalized to the length of the worm, which is outputted as TOF (time of flight) from the Biosorter. At least 3 biological replicates were completed per condition, with a minimum of 100 animals per replicate. Statistical analysis was done using the student’s t-test analysis in GraphPad Prism version 10.0.0 for Windows, GraphPad Software, Boston, Massachusetts USA, www.graphpad.com.

### Drug Screen and UPR^mt^ Assessment via CX5

First, large number of embryos were harvested via bleaching and plated on NGM plates with OP50. 3 days later, 0.7 OD of OP50 bacteria in S basal medium was used to wash off the worms from the NGM plates and ∼ 50 worms were added per well of a 384-well plate (ThermoFisherScientific, 142761). Subsequently, treatment solutions were added to a total of 50 µl solution with 10 wells per condition, and then the plate was incubated at 20°C for 24 hours. Then 50 µl of 0.04% Sodium Azide (VWR, 26628-22-8) was added to each well to paralyze the worms. The 384-well plate was scanned using the CellInsight CX5 HCS Platform (ThermofisherScientific, CX51110) at a light intensity of 90%, an exposure of 40%, and a pixel intensity threshold of 2000 (**Fig 3A**).

Visual inspection of the images was carried out to censor wells that were out of focus, had debris in it, or had too few worms. The “spottotalintench3” data, green fluorescence quantifying mitochondrial stress, was then normalized to “spotcountch2” data, red fluorescence used to count worm number, per well to graph mitochondrial stress. At least 3 biological replicates were completed per condition, each condition contained 10 wells with between 30-50 animals in each well. Statistical analysis was done using the student’s t-test analysis in GraphPad Prism version 10.0.0 for Windows, GraphPad Software, Boston, Massachusetts USA, www.graphpad.com.

### Competition Assay

Three biological replicates of the competition assay were done on OP50 seeded NGM plates with 7 N2 (wildtype) animals and 7 *uaDf5* animals and the control “non-competition” plates had 14 *uaDf5* animals to start. These plated were passaged bi-weekly from the previous generation’s plates using the same ratio of animals. Assay was continued until *uaDf5* animals on the “competition” plate were outcompeted by the N2 animals.

### Animal Length Assay

Thirty L4 animals were picked per replicate, where 10 animals were imaged at L4, at Day 2, and Day 4 of adulthood. On the day of imaging, 10 animals per genotype were picked onto NGM plates without bacteria and paralyzed with 10% Sodium Azide. Images were taken on a Stemi 305 Zeiss Microscope, and Image J^32–34^ was used to measure the length of individual worms. Statistical analysis was done using the student’s t-test analysis in GraphPad Prism version 10.0.0 for Windows, GraphPad Software, Boston, Massachusetts USA, www.graphpad.com.

### Lifespan Analysis

For each strain/condition, 3 replicates of 30 L4 animals were grown at 20°C, every other day the worms were transferred onto fresh plates (with treatment administered as described above). When worms were transferred onto fresh plates they were categorized as either alive, dead, or censored. If a worm was not moving at the time of transfer, they were gently prodded and counted as dead if there was no response. Censored animals, worms that escaped or died aberrantly, such as bagging or having a burst vulva, were removed from final analysis. Analysis was completed using GraphPad Prism version 10.0.0 for Windows, GraphPad Software, Boston, Massachusetts USA, www.graphpad.com.

## RESULTS

### Fitness defects severity correlated with increasing *uaDf5* heteroplasmy levels

The *uaDf5* animals carry a 3,053 bp heteroplasmic mtDNA large-scale deletion encompassing 4 mRNA and 7 tRNA coding genes (**Fig 1A, B**). Three separate sets of *uaDf5* strains were studied: the *uaDf5* strain without any reporters, one set of strains with the *hsp-6p::GFP* reporter constructs (low and medium heteroplasmy), and one set of strains carrying the *hsp-6p::GFP and myo-2p::mCherry* reporters (low, medium, and high heteroplasmy) (**Supp. Fig 1A**). The strain without any reporters had an average of 2.3% heteroplasmy with a range from 0.4% to 5.6% heteroplasmy (**Supp. Fig 1B, C**), and no fecundity defect (**Supp. Fig 1D**). The low heteroplasmy *uaDf5* (#1) strain carrying *hsp-6p::GFP* showed no change in total mtDNA content (**Fig 1C**) as compared to wild-type (WT) animals (*hsp-6p::GFP*) and exhibited an average of 18.6% heteroplasmy (p<6.9×10^-4^) (**Fig 1C, D**). The medium heteroplasmy *uaDf5* (#2) strain carrying *hsp-6p::GFP* showed a 1.8-fold increase (p<0.016) in total mtDNA content (**Fig 1C**) compared to WT (*hsp-6p::GFP*) and exhibited an average of 35.5% (p<3.8×10^-7^) heteroplasmy (**Fig 1C, D & Supp. Fig 1E**); heteroplasmy levels increased from an average of 13.9% at larval stage 4 (L4) to an average of 28% at Day 10 of adulthood (**Fig 1E**). The low and medium heteroplasmy *uaDf5* strains carrying *hsp-6p::GFP* showed increasing levels of 1.5-fold (p<0.039) and 3.3-fold (p<0.0018) higher than WT levels of mitochondrial unfolded protein response (UPR^mt^) induction, which was assessed by quantifying *hsp-6p::GFP* fluorescence normalized to worm length on the COPAS Biosorter (**Fig 1F & Supp. Fig 1F**). Both *uaDf5* strains carrying *hsp-6p::GFP* showed a minimal (8%) decrease in progeny count (p<0.023, p<0.044) (**Fig 1G**) with no defect in thrashing abilities at day 5 (**Fig 1H**) or day 7 (**Supp. Fig 1G**) of adulthood compared to WT (*hsp-6p::GFP*).

**Figure 1:**
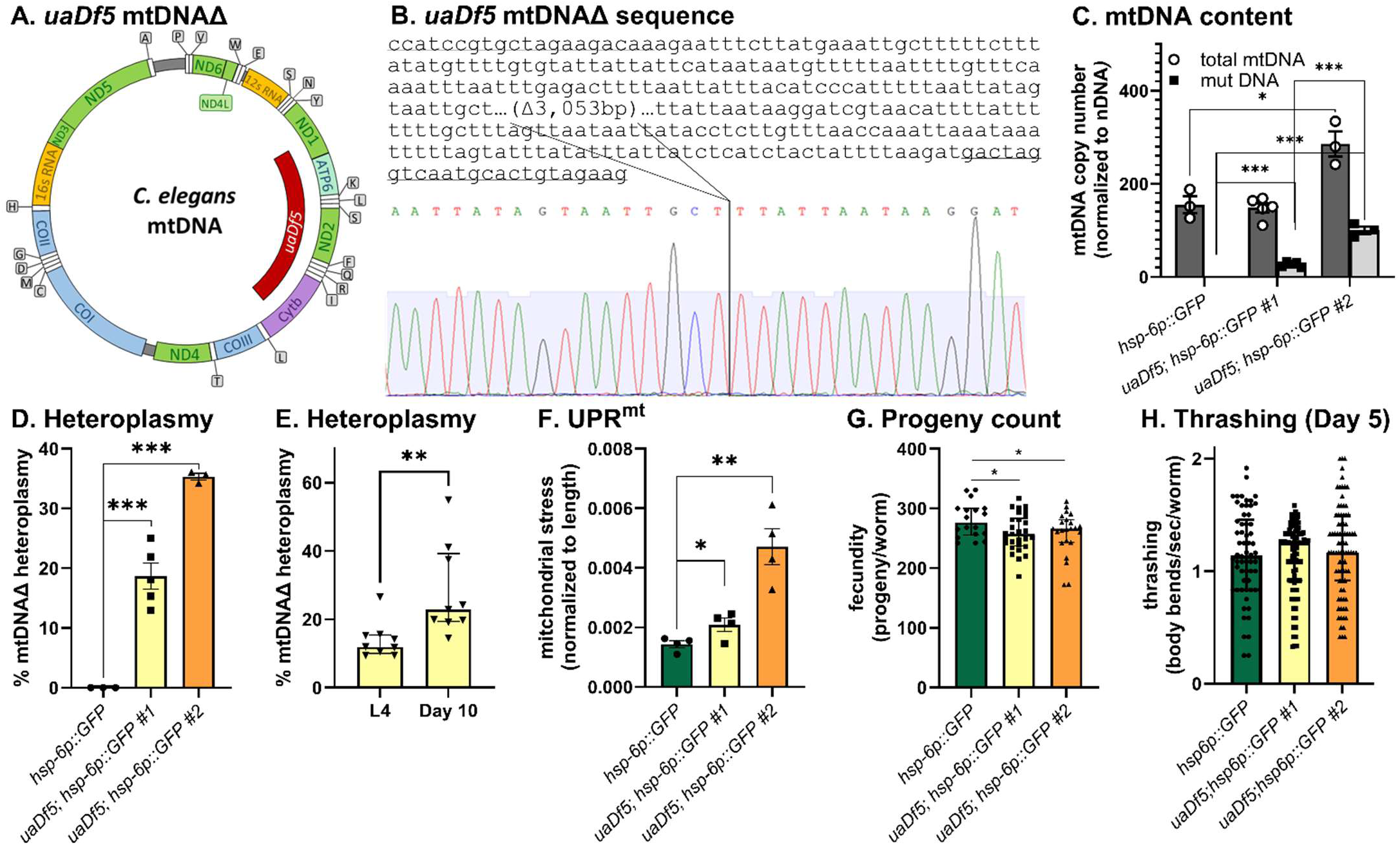
Up to 35% heteroplasmy in *uaDf5* strains gives rise to mild phenotype. **(A)** *C. elegans* mtDNA highlighting protein (complex I (green), complex III (purple), complex IV (blue), complex V (light green)), rRNA (orange) and tRNA (lettered boxes) coding genes. The inner bar (red) represents the ∼3.1kb deletion in the *uaDf5* animals that impacts 4 protein-coding and 7 tRNA genes. **(B)** Partial *C.elegans* mtDNA sequence and an electrophoretogram depicting the *uaDf5* deletion breakpoints. **(C)** Total mtDNA (dark gray) was 1.8x higher (p<0.016) in *uaDF5;hsp-6p::GFP #2* compared to *hsp-6::GFP*. The *uaDF5;hsp-6p::GFP #1* and *uaDf5;hsp-6p::GFP #2* strains had an average of 27 (p<1.2×10^-6^) and 100 (p<1.2×10^-6^) copies of mutant mtDNA per nuclear genome. **(D)** On average, the *uaDF5;hsp-6p::GFP #1* (yellow) and *uaDf5;hsp-6p::GFP #2* (orange) strains exhibited 18.6% (p<6.9×10^-4^) and 35.3% (p<3.8×10^-7^) heteroplasmy levels. **(E)** At day 10 of age, *uaDf5;hsp-6p::GFP #1* exhibited an average of 28% heteroplasmy, a 2x increase (p<0.0086) compared to their siblings at L4 stage that exhibited an average of 13.9% heteroplasmy. **(F)** The *uaDF5;hsp-6p::GFP #1* (yellow) and *uaDf5;hsp-6p::GFP #2* (orange) strains exhibited 1.5x (p<0.039) and 3.3x (p<0.0018) more UPR^mt^ relative to *hsp-6p::GFP* (green) strain when assessed on the COPAS biosorter. **(G)** Both the *uaDF5;hsp-6p::GFP #1* (yellow) and *uaDf5;hsp-6p::GFP #2* (orange) strains exhibited an 8% decrease in progeny count (p<0.023, p<0.044) compared to *hsp-6p::GFP* (green) animals. **(H)** Neuromuscular fitness assessed via thrashing, where the data represents body bends per second per worm, showed no defect in the *uaDF5;hsp-6p::GFP #1* (yellow) and *uaDf5;hsp-6p::GFP #2* (orange) strains as compared to *hsp-6p::GFP* (green) animals at Day 5 of adulthood. For figures (C)-(E), each data point represents a biological replicate, at least 3 biological replicates were done, and the mean±SEM is graphed. At least 3 separate biological replicates were completed, where the data per worm is graphed for figure (G)(H) and the data per sample of 30 worms was graphed for figure (H) with the median+interquartile range. Student’s t-test analysis was performed to assess significance for figures (C)-(H).

In generating the *uaDf5* strain with the *hsp-6p::GFP* and *myo-2p::mCherry* reporters to facilitate higher throughput screening capabilities on a high content imager, we isolated 3 different strains with low, medium and high heteroplasmy (Low Δ %, Med Δ %, High Δ %) strains (**Supp. Fig 1A**) that exhibited on average 13.9% (p<0.00017), 29.3% (p<1.1×10^-6^) and 62% (p<2.6×10^-4^) heteroplasmy (**Fig 2A, B**). The Low Δ % and Med Δ % strains had a 1.4x (p<0.013) and a 1.6x (p<0.0037) increase in total mtDNA content compared to WT (*myo-2p::mCherry;hsp-6p::GFP*) (**Fig 2A**). The Low, Med, and High heteroplasmy strain exhibited increasing levels of 97-fold (p<0.015), 223-fold (p<1.8×10^-5^), and 483-fold (p<2.1×10^-6^) higher induction relative to WT UPR^mt^ as quantified by *hsp-6p::GFP* fluorescence normalized to worm count per well on the high content imager CX5 (**Fig 2C & Supp. Fig 2A**). UPR^mt^ analysis on the COPAS Biosorter in the High Δ % animals showed an increase with age over a 98-hour period (**Supp. Fig 2B**). At 98-hours, those with <750 time of flight (larval stage animals) had maximal UPR^mt^ induction while those animals with >750 time of flight (adult stage) already started exhibiting lower UPR^mt^ (**Supp. Fig 2C**). The High Δ % strain exhibited a developmental delay where after 4 days of growth from embryos, 16.6% of the population were young adults compared to 1.5% of the population for WT (p<0.0036) (**Fig 2D**). The High Δ % strain exhibited a 17.8% size defect when assessed on the COPAS Biosorter compared to WT (green) (p<0.013) (**Fig. 2E**), which arose by 48-hours post harvesting embryos and almost completely disappeared by 98-hours (**Supp. Fig 2D**). Synchronizing the worms at L4 stage and imaging showed a 6% size increase at L4 stage (p<0.01) and 6.7% size defect at Day 2 of adulthood (p<6.1×10^-4^), which disappeared by Day 4 of adulthood (**Fig 2F & Supp. Fig 2E**). These observations track with the delayed development phenotype in **Fig 2D**, which accounts for most of the length defect observed in the COPAS Biosorter experiment in **Fig 2E** which disappears over time (**Fig 2F & Fig Supp. Fig 2D, E**). The High Δ % strain exhibited 88.4% (p<3.4×10^-15^) reduced progeny count (**Fig. 2H**) but showed no defect in the hatch rate (**Fig. 2G**) compared to the WT. The High Δ % strain exhibited 50% (p<0.038) reduced thrashing rate at L4 relative to the WT animals (**Fig. 2I**). The High Δ % strain exhibited no lifespan defects (**Fig 2J**), however, 28.7% of the population died from aberrant deaths such as bagging and burst vulva compared to 0.88% of the population in WT (**Fig 2K**). In competitive growth with WT animals, the High Δ % animals (brown) are outcompeted by the 5^th^ generation (**Fig. 2L**).

**Figure 2:**
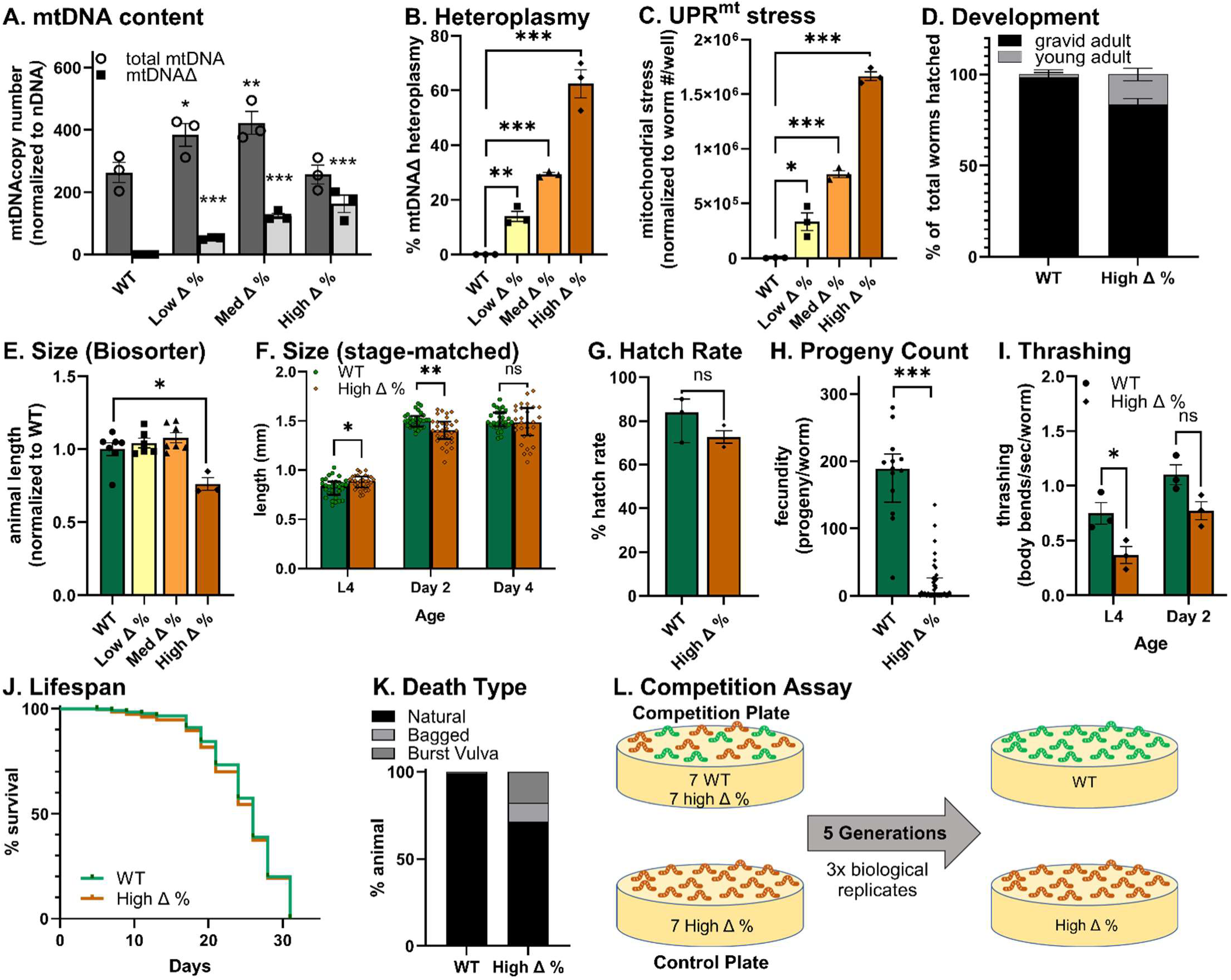
Greater than 50% heteroplasmy in *uaDf5* strains gives rise to a more severe phenotype. **(A)** The Low Δ %, Med Δ %, High Δ % strains respectively had a total mtDNA copy number (dark gray) of 384, 422, and 257 per haploid genome, which was a 1.4x (p<0.013) and a 1.6x (p<0.0037) increase in the Low Δ % and Med Δ % strains compared to the 263 total mtDNA copy number per haploid genome observed in WT animals. The Low Δ %, Med Δ %, High Δ % strains had an average of 52 (p<1.6×10^-5^), 123 (p<1.2×10^-4^), and 162 (p<0.0043) mutant mtDNA (light gray) per haploid genome. **(B)** On average, the Low Δ % (yellow), Med Δ % (orange) and High Δ % (brown) strains respectively exhibited 13.9% (p<0.00017), 29.3% (p<1.1×10^-6^) and 62% (p<2.6×10^-4^) heteroplasmy. **(C)** The Low Δ % (yellow), Med Δ % (orange) and High Δ % (brown) strains exhibited 97x (p<0.015), 223x (p<1.8×10^-5^), and 483x (p<2.1×10^-6^) more UPR^mt^ relative to the wildtype (WT - green) strain when assessed on the high content imager CX5. **(D)** The High Δ % (brown) strain exhibited a developmental delay where after 4 days of growth from embryos, 16.6% of the population were assessed as young adults compared to 1.5% of the population for WT animals (p<0.0036). **(E)** The High Δ % (brown) strain, synchronized via bleaching, exhibited a 17.8% size defect (p<0.013) when assessed on the COPAS biosorter compared to WT (green) animals. **(F)** The High Δ % (brown) strain, synchronized by picking L4 animals, exhibited a 6% size increase at L4 stage (p<0.01) and 6.7% size defect at Day 2 of adulthood (p<6.1×10^-4^) compared to WT (green) animals. **(G)** The High Δ % (brown) strain exhibited no defect in the hatch rate compared to WT (green) animals. **(H)** The High Δ % (brown) strain exhibited 88.4% reduction (p<3.4×10^-15^) in progeny count relative to WT (green) animals. **(I)** The High Δ % (brown) strain exhibited 50% reduced thrashing rate at L4 (p<0.038) and 30% reduced thrashing at Day 2 of adulthood (ns = p<0.055) relative to WT (green). **(J)** The High Δ % strain (brown line) exhibited no lifespan defects compared to WT (green line) animals. However, **(K)** 28.7% of the population died from aberrant deaths such as bagging and burst vulva compared to 0.88% of the population in WT animals. **(L)** In competitive growth with WT animals (green), the High Δ % animals (brown) were outcompeted by the WT population by the 5^th^ generation. Each data point represents a biological replicate and at least 3 biological replicates were done for figures (A-I)(K), with the mean±SEM graphed for figures (A-C)(E)(G)(I) and the data per worm is represented with the median+interquartile range in for (F-H). Student’s t-test analysis was performed to assess significance for (A-I) and the Gehan-Breslow-Wilcoxon test was performed to assess significance for figure (J).

**Figure 3:**
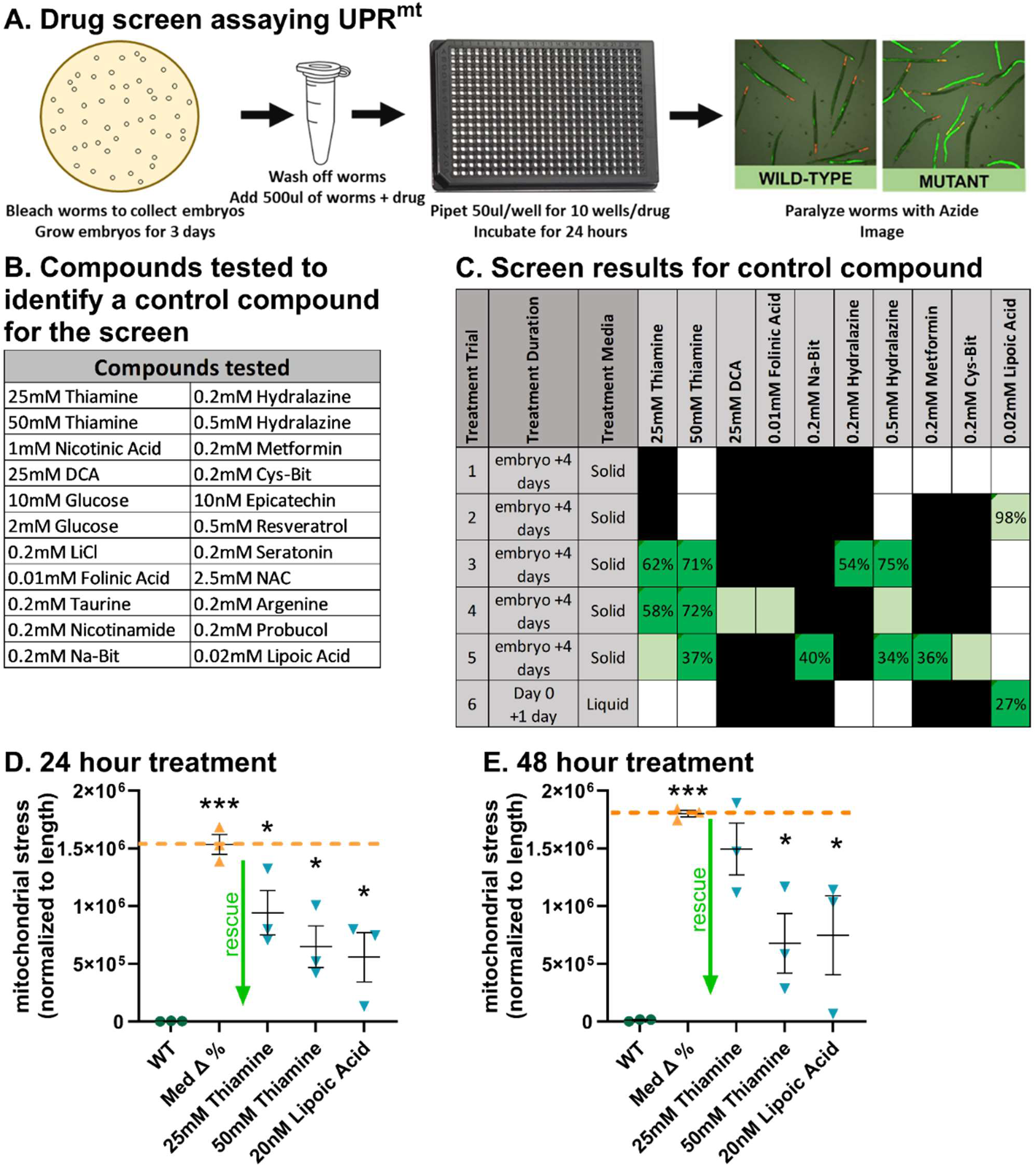
Identifying a control treatment to utilize for the mitochondrial stress or high-throughput screen. **(A)** Schematic overview of the screening protocol assaying UPR^mt^ where worms were synchronized by collecting embryos on day 0 and grown on NGM plates with OP50 for 3 days. On day 3, the worms were collected in 500µl S-Basal Media + treatment and aliquoted at 50µl/well to be incubated at 20°C for 24 hours. On day 4 the worms were imaged on the high-content imager CX5. **(B)** List of compounds and the concentrations at which they were tested at least once in 6 trials to identify a positive control condition for use in the screen of mitophagy modulators. **(C)** Table summarizing the screen results for the positive controls identified to reliably reduces UPR^mt^. The drug exposure spanned from embryogenesis + 4 days on NGM plates with OP50 for trials 1-5 and L4 stage+1 day in S basal medium with OP50 for trial 6. The white boxes indicate untested drug in that trial, black boxes indicate no rescue, light green indicate a rescue trend without statistical significance, dark green boxes indicate statistically significant rescue and the % rescue is numerically indicated. **(D)** L4+1 day in S basal medium with OP50 showed an average of 39% reduction with 25mM Thiamine (blue, p<0.048), 58% reduction with 50mM Thiamine (blue, p<0.011), and 64% reduction with 20nM Lipoic Acid (blue, p<0.013) in mitochondrial stress compared to untreated Med Δ % animals (orange). **(E)** L4+2 days treatment in S-medium with OP50 showed an average of 62% reduction with 50mM Thiamine (blue, p<0.013), and 59% reduction with 20nM Lipoic Acid (blue, p<0.037) in mitochondrial stress as compared to untreated Med Δ % animals (orange). For (D, E), each data point represents a biological replicate, 3 biological replicates were performed, and the mean±SEM is graphed. Student’s t-test analysis was performed to assess significance.

The High Δ % strain, when maintained in crowded but not starved conditions at 20°C, had reduced heteroplasmy by 32% (p<7.7×10^-4^) over 1 month (∼8 generations) and 40% (p<0.0093) over two months (∼16 generations) (**Supp. Fig 2F, G**). The animals with lower heteroplasmy after one month also showed improvement in fitness defects, such as the loss of developmental delay (**Supp. Fig 2H**), 2-fold increase in fecundity (**Supp. Fig 2I**), and WT levels of movement (**Supp. Fig 2J**).

### Mitophagy modulating compound screening in the Med Δ % ***uaDf5*** by assay of UPR^mt^ induction

The UPR^mt^ was assessed using the CX5, as described above, and the few parameters that needed troubleshooting were (i) the use of Spot Intensity versus Total Intensity to measure the UPR^mt^, (ii) threshold setting prior to imaging, and (iii) duration of compound exposure. The Spot Intensity (**Supp. Fig 3A**) versus the Total Intensity (**Supp. Fig 3B**) outputs were tested and the Spot Intensity output (**Supp. Fig 3A**) clearly reported a 5.8-fold (p<2.3×10^-8^) higher mitochondrial stress observed visually (**Supp. Fig 2A**) in Med Δ % animals compared to WT animals. Testing threshold levels assessing a globally fluorescing mitochondrial worm (*cox-4(zu476[cox-4::eGFP::3xFLAG)*) showed a linear correlation in all samples between number of worms and the total fluorescence at 40% exposure (**Supp. Fig 3D**) but not at 25% exposure (**Supp. Fig 3C**). Twenty-four compounds empirically observed to rescue fitness defect in the mitochondrial complex I mutant *gas-1(fc21)* strain were tested in the Med Δ % to identify a positive control compound that can reproducibly reduce UPR^mt^ (improve mitochondrial health) for use in the mitophagy modulating drug screen (**Fig 3B**). While testing the 24 compounds, long treatment of 4-day exposure on solid media (embryogenesis to Day 1 of adulthood) and a short treatment of 24-hour exposure in liquid media (L4 to Day 1 of adulthood) were tested (**Fig 3C**). Thiamine showed strong reproducible effects and lipoic acid showed a potential for very strong rescue. They were tested in multiple biological replicates for 24-hour (**Fig 3D**, **Supp. Fig 3E-G**) and 48-hour (**Fig 3E**, **Supp. Fig 3H-I**) liquid exposure starting at L4 stage. All three treatments (25 mM Thiamine, 50 mM Thiamine and 20 nM Lipoic acid) showed significant and reproducible (39% (p<0.048), 58% (p<0.011), and 64% (p<0.013)) UPR^mt^ reduction at 24-hour exposure, which was the duration of treatment condition used in the screen of the 62-mitophagy modulating drugs (**Fig 3A**).

The 62-mitophagy modulating compound library was tested twice at 25 µM, where ∼20 compounds were tested per plate. The control wells (WT, Med Δ %, 0.13% DMSO and 50 mM Thiamine) were combined across the 3 plates per screen replicate and the data was displayed in two graphs (**Supp. Fig 4**). Analysis of the first replicate, in the comparisons using combined controls, identified compounds URB-597, Hemin, Celastrol, Esmolol (hydrochloride), and Clioquinone to reduce UPR^mt^ by 28% (p<0.046), 50% (p<1.9×10^-4^), 25% (p<0.012), 30% (p<0.020), and 28% (p<0.038) respectively (**Supp. Fig 4A &B**). Identical analysis of the second replicate, in the comparisons using combined controls, identified compounds Hemin, Resveratrol, and Pifithrin-α (hydrobromide) treatment to reduce UPR^mt^ by 33% (p<0.0017), 38% (p<0.0036), and 20% (p<0.038) respectively (**Supp. Fig 4C & D**). The screen findings are summarized in **Supp. Fig 5**.

Compound treated Med Δ %, compared to solvent treatment (DMSO or Water) on the same plate identified Vorinostat, Hemin, GSK2578215A, Celastrol and Resveratrol to reduce UPR^mt^, an analysis we found more reliable. These five compounds tested at 4 different doses (12.5 µM, 25 µM, 50 µM and 100 µM) showed that only Hemin and Celastrol reduced UPR^mt^ in a dose-dependent manner (**Fig 4A & B**). Six replicate treatments with 12.5µM, 25µM, 50µM and 100µM of Hemin reproducibly reduced UPR^mt^ by 26% (p<0.012), 46% (p<0.012), 69% (p<0.00132) and 72% (1.0×10^-5^) and treatments with 12.5 µM, 25 µM, 50 µM and 100 µM Celastrol reproducibly reduced UPR^mt^ by 22% (non significant), 37% (p<0.0044), 62% (p<1.2×10^-4^) and 56% (p<0.0085). 100 µM Hemin and Celastrol treatments result in dead animals. Combinatorial testing of Hemin and Thiamine or Celastrol and Thiamine showed no additive synergistic effects. In fact, 12.5 μM Hemin + 10 mM Thiamine co-treatment increased UPR^mt^ by 1.5-fold (p<0.0017) compared to 12.5 μM Hemin treatment alone (**Fig 4C**). 25 μM Celastrol + 10 mM Thiamine co-treatment increased UPR^mt^ 1.3-fold (p<0.032) compared to 25 μM Celastrol treatment alone (**Fig 4C**). 12.5 μM Celastrol + 25 mM Thiamine co-treatment increased UPR^mt^ by 1.4-fold (p<0.0068) compared to 25 mM Thiamine treatment alone (**Fig 4C**). To study the impact of Hemin on additional fitness outcomes, 2 batches of Hemin was obtained from MedChemExpress, one with the same batch number of Hemin in the library and the second one had a different batch number. Neither batch of Hemin reduced UPR^mt^ at 25 µM, 50 µM, and 100 µM (**Supp. Fig 6A**) and one batch was able to reduce UPR^mt^ by only 45% (p<1.7×10^-9^) at 150 µM. The other batch showed an increase in UPR^mt^ by 1.3x (p<0.049) at 100 µM. Even after sonication, which should help solubilize compounds, there was no reduction in UPR^mt^ (**Supp. Fig 6A**). Hemin sourced from Cayman Chemical showed reduction in UPR^mt^ at the highest dose of 200 µM but was not as effective as the Hemin obtained from the MedChemExpress library (**Supp Fig 6C**). Further studies with Hemin were halted due to the unreliable and low efficacy of the compound. The batch of Celastrol obtained from MedChemExpress also failed to reduce UPR^mt^ at 25 µM, 50 µM, 100 µM and 150 µM (**Supp. Fig 6B**). Testing Celastrol from Selleck Chemicals, Sigma-Aldrich and Cayman Chemical showed that the Cayman Celastrol performed the best of the three sources, which at 50 µM, 100 µM and 150 µM, reduced UPR^mt^ by 58% (p<7.4×10^-6^), 62% (p<1.6×10^-6^), and 63% (p<1.4×10^-6^), with no deaths at any concentration (**Supp Fig 6C**); therefore, this formulation was used for remainder of the studies.

**Figure 4:**
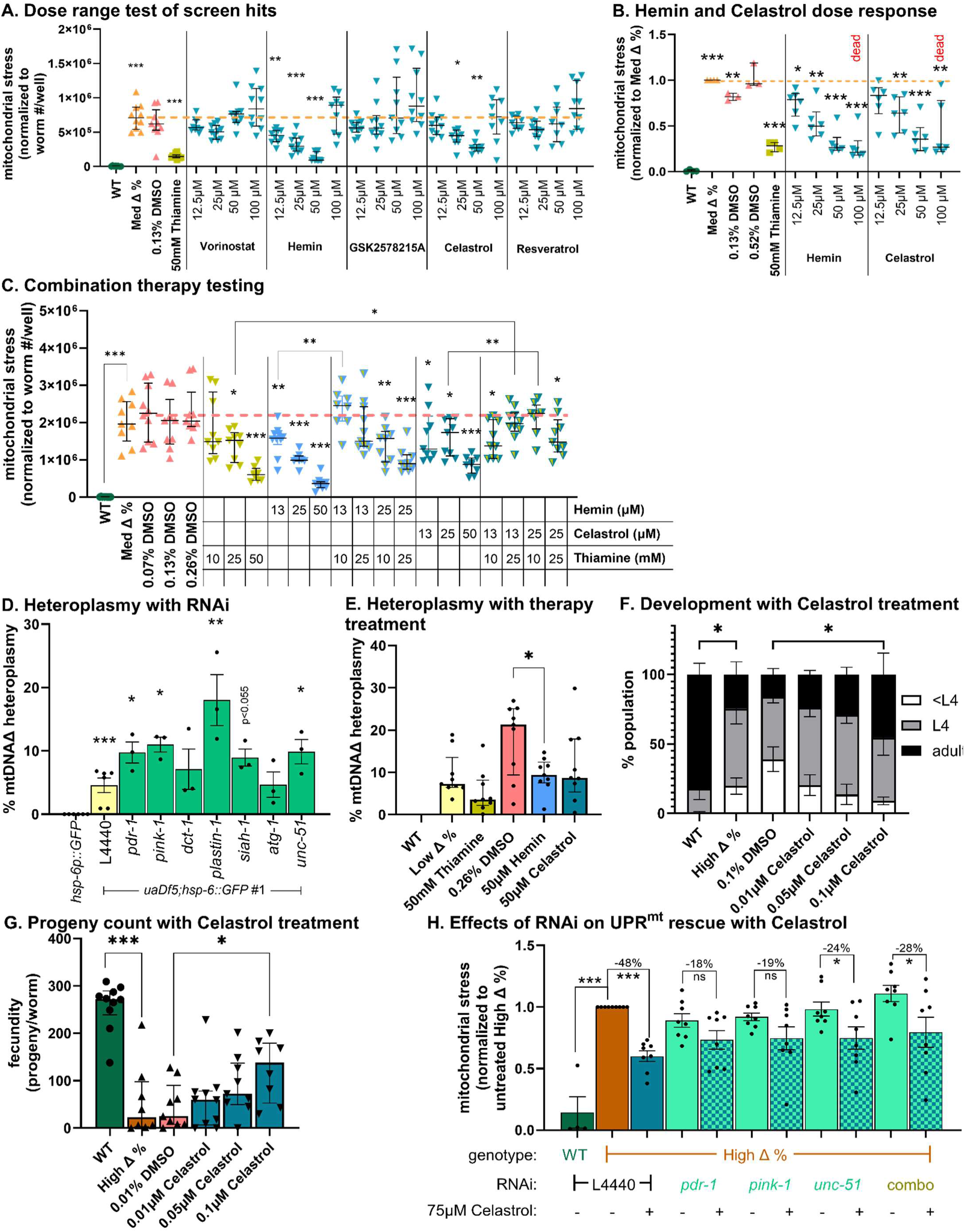
Celastrol rescues disease phenotypes in *uaDf5* animals partly via mitophagy modulation. **(A)** Treatment of Med % Δ with 4 different doses (12.5µM, 25µM, 50µM and 100µM) of the 5 hits (Vorinostat, Hemin, GSK2578215A, Celastrol and Resveratrol) (blue), showed that 12.5 µM (p<0.0033), 25µM (p<6.4×10^-5^) and 50µM (p<2.2×10^-8^) Hemin, and 25µM (p<0.0034) and 50µM (p<1.8×10^-5^) Celastrol reduced mitochondrial stress in a dose-dependent manner. **(B)** Six replicate treatments of Med Δ % animals with 12.5µM, 25µM, 50µM and 100µM of Hemin reduced UPR^mt^ by 26% (p<0.012), 46% (p<0.012), 69% (p<0.00132) and 72% (1.0×10^-5^) and of Celastrol reduced UPR^mt^ by 22% (n.s.), 37% (p<0.0044), 62% (p<1.2×10^-4^) and 56% (p<0.0085). 100µM Hemin and Celastrol treatments resulted in dead animals. **(C)** Individual treatment with (25mM (p<0.18) and 50mM (p<1.3×10^-5^)) Thiamine (light green), (13µM (p<0.008), 25µM (p<2.6×10^-5^), and 50µM (p<6.5×10^-10^)) Hemin (blue), and (13µM (p<0.042), 25µM (p<0.023), and 50µM (p<1.1×10^-6^)) Celastrol (dark blue) reduces UPR^mt^. Co-treatment of Thiamine with Hemin or Celastrol did not result in synergistic reduction of UPR^mt^. 12.5μM Hemin + 10mM Thiamine co-treatment increased UPR^mt^ 1.5-fold (p<0.0017) compared to 12.5μM Hemin treatment alone. 25μM Celastrol + 10mM Thiamine co-treatment increased UPR^mt^ 1.3-fold (p<0.032) compared to 25μM Celastrol treatment alone. 12.5μM Celastrol + 25mM Thiamine co-treatment increased UPR^mt^ by 1.4-fold (p<0.0068) compared to 25mM Thiamine treatment alone. **(D)** RNAi against mitophagy genes showed that *pdr-1*, *pink-1*, *plastin-1*, and *unc-51* knockdown (green) increased the heteroplasmy by 2.1x (p<0.037), 2.2x (p<0.010), 3.0x (p<0.0035), and 2.1x (p<0.040) compared to the low heteroplasmy *uaDf5;hsp-6::GFP* #1 strain (yellow). **(E)** Heteroplasmy analysis showed that 50µM Hemin treatment (blue) from embryogenesis + 4 days resulted in a 47% decrease in heteroplasmy (p<0.016) as compared to 0.26% DMSO treated Med Δ % animals (yellow). 50mM Thiamine (light green) and 50µM Celastrol (dark blue) treatment showed non-significant trends of lowering the heteroplasmy by 43% and 36%, respectively, as compared to untreated and 0.26% DMSO treated Med Δ % animals. **(F)** Assaying development at embryo + 3 days showed significant delay in High Δ % animals compared to WT, when on average the High Δ % animals are 20% <L4 (p<0.0.31), 56% L4 (p<0.047) and 25% adults (p<.0097) compared to 0.7% <L4, 17% L4 and 82% adults in WT animals. Treatment with 0.1µM Celastrol rescued development rate with an average of 9%<L4 (p<0.31), 46% L4 and 25% adults 3 days post embryogenesis compared to the DMSO treated animals, which were at an average of 40%<L4 (p<0.31), 44% L4 and 16% adults. **(G)** The High Δ % (brown) strain exhibited 78% reduction (p<4.4×10^-6^) in progeny count relative to WT (green) animals. Treatment with 0.1µM Celastrol (blue) improved progeny count by 2.6-fold (p<0.015) compared to DMSO treated (pink) animals. **(H)** The efficacy of 75µM Celastrol is diminished to −17% (n.s), −19% (n.s.), −23% (p<0.046), and −28% (p<0.039) in the presence of RNAi knockdown of the mitophagy genes *pdr-1*, *pink-1*, *unc-51*, and combination of all three RNAi (light green and checkered) as compared to the −47% (p<6.8×10^-8^) reduction of UPR^mt^ conferred by just Celastrol alone (dark blue) in High Δ % animals. For (A)(C), each data point represents a technical replicate, 8-11 technical replicate was acquired for each condition, and the median+interquartile range is graphed. For (B)(D)(E)(G)(H), each data point represents a biological replicate, 3-9 biological replicates were performed. The mean±SEM is graphed for (B)(H) and the median±interquartile range is graphed for (E)(G). For (F), the data represents the combination of 3 biological replicate experiments with at least 30 animals per condition. Student’s t-test analysis was performed to assess significance. (n.s.) applies to no statistical significance.

### Celastrol rescued multiple disease outcomes and functions partly via mitophagy regulation

We observed that the individual RNAi knockdown of mitophagy genes^36–38^, *pdr-1*, *pink-1*, *plastin-1*, *siah-1,* and *unc-51*, increased the heteroplasmy by 2.1-fold (p<0.037), 2.2-fold (p<0.010), 3.0-fold (p<0.0035), 2.0-fold (non-significant), and 2.1-fold (p<0.040), respectively, as compared to the control L4440-fed animals (**Fig 4D**). Treatment with the mitophagy activator 50 µM Hemin (sourced from the mitophagy modulating drug library) showed a 47% (p<0.016) decrease in heteroplasmy (**Fig 4E**). Treatment with 50 mM Thiamine and 50 µM Celastrol, a mitophagy activator, showed non-significant trends of lowering the heteroplasmy by 43% and 36% (**Fig 4E**). Treatment with 0.1 µM Celastrol rescued development defect, where there was a 75% reduction in the young larva population (<L4) compared to solvent treated High Δ % animals (**Fig 4F**). Treatment with 0.1 µM Celastrol also improved fecundity, where a 2.6-fold (p<0.015) increase in progeny count was observed when compared to solvent treated High Δ % animals (**Fig 4G**). The observed 47% (p<6.8×10^-8^) reduction in mitochondrial with 75 µM Celastrol treatment was diminished to non-significant change in UPR^mt^ upon the downregulation of either *pdr-1* or *unc-51* and down to 23% (p<0.046) upon downregulation of *unc-51* (**Fig 4H**). Simultaneous knock down of all three mitophagy components (*pdr-1*, *pink-1*, *unc-51*) reduced the Celastrol mediated decrease in mitochondrial stress induction to 28% (p<0.039) (**Fig 4H**).

## DISCUSSION

*uaDf5* strains harboring variable SLSMD heteroplasmy levels recapitulated salient features of human SLSMD disease. Most broadly, we found that while sibling worms each displayed ranging levels of heteroplasmy, the average population heteroplasmy levels remained similar over multiple generations. Variable heteroplasmy levels amongst siblings directly mimics what is observed in human pedigrees diseases arising from maternally inherited mtDNA mutations. This is generally due to the germline bottleneck and stochastic uneven distribution of mtDNA that occurs during cell division and oogenesis^39–41^.

The *uaDf5* heteroplasmy levels strongly correlated with the severity of fitness defects. Low, middle and high heteroplasmy levels of the mtDNA SLSMD led to increasing levels of UPR^mt^ induction, as quantitatively assessed by a fluorescent reporter strain. This direct correlation between the increasing level of heteroplasmy and increasing phenotypic severity aligns with expectations. Furthermore, the higher heteroplasmy strain exhibited additional loss in organismal fitness phenotypes beyond mitochondrial stress induction, manifesting developmental, fecundity, and neuromuscular (thrashing) defects. The SLSMD heteroplasmy level at which point these additional features started exhibiting clearly represents an important threshold where the dysfunction we observe at the cellular level (UPR^mt^), which likely correlated with worsening mitochondrial respiratory function, becomes sufficiently severe that the mitochondrial dysfunction manifests at the organismal level with reduced fitness. These findings also closely align with the standard paradigm for biochemical or clinical threshold that is clinically relevant, as it represents a level of heteroplasmy that gives rise to clinical symptoms. Interestingly, the SLSMD heteroplasmy levels fell over generations, where in the higher heteroplasmy *uaDf5* lines we observed a direct correlation between decreasing heteroplasmy levels and phenotypic severity. This observation cements the finding that higher heteroplasmy causes severe fitness defects, rather than these phenotypes being an artifact of a spontaneous background mutation. This is consistent with studies in mice that suggest there is a minimum percentage of healthy mtDNA genomes necessary to maintain adequate mitochondrial respiratory chain function, taking into account both heteroplasmy level and overall mtDNA content, underlying the physiologic effects of rising levels of mtDNA heteroplasmic mutations^42^. Interestingly, in some mtDNA disorders, it has been well noted that heteroplasmy levels may fall over time in rapidly dividing tissues (such as blood) even when they rise with age in affected tissues^43^.

The phenotype in *uaDf5* animals correlate with SLSMD disease symptoms. As expected, the *uaDf5* animals have quantifiable levels of SLSMD heteroplasmy ranging from 10% to 70%, directly correlating with their respective levels of mitochondrial stress levels. In particular, significant neuromuscular defects occurred in the *uaDf5* animals in the form of reduced body movement (thrashing) and an increase in aberrant deaths. While the connection between body movement defects and neuromuscular dysfunction is apparent, the types of aberrant death we observe (bagging, a result of embryo laying defect and exploding vulva) are also indicative of a vulval defect, which is neuromuscular by nature. These are important observations since SLSMD patients largely have severe neuromuscular weakness. Ultimately, identification of treatments that improve neuromuscular defects in *uaDf5* animals will hold the potential for neuromuscular function improvement in SLSMD patients.

Several strains have now been identified in *C. elegans* that carry a range of different size mtDNA deletions. These can be further developed to gain deeper understanding into SLSMD pathogenesis and to preclinically evaluate therapeutic strategies. In future studies, it would be beneficial to introduce in *C. elegans* similar strategies for mtDNA engineering as are being developed in vertebrate system. For example, we postulate that a genetically-encoded constitutively active mtDNA editor may offer one strategy to combat the SLSMD heteroplasmy loss that occurs over generations to generate a stable heteroplasmic model of SLSMD disease. Such strategies will expand the genetic toolkit we have available to model SLSMDS, as a similar inability to maintain heteroplasmic SLSMD mutations has long been reported in mammalian models^44^. Increasing mitophagy is likely to promote selective degradation of dysfunctional mitochondria, which can reasonably be presumed to carry higher loads of mutant mtDNA genomes. Thus, mitophagy would be expected to induce a preferential degradation of mutant mtDNA genomes, thereby reducing the mutant heteroplasmy levels and resulting cellular pathophysiology that causes multi-systemic disease phenotypes. Indeed, previous studies have shown that genetic perturbations to mitophagy do modulate heteroplasmy levels, similarly as we also observed using RNAi knockdown of key mitophagy genes’ expression^21,25^. These data support the rationale to develop mitophagy modulators as therapies for SLSMD disorders while directly target the etiology of the disease. We successfully employed this strategy to identify two mitophagy activators, Hemin^45–47^ and Celastrol^48^. Hemin is an erythroid maturation stimulator and has been reported to activate mitophagy in leukemic erythroblast cell line^49^. It has been historically developed as a treatment for acute attacks of porphyria^45^ and more recently in inflammatory diseases^47^. Currently it is being repurposed for potentially treating cardiovascular diseases^46^. Celastrol is mitophagy activating anti-inflammatory, anti-cancer, and metabolic modifying natural product derived compound and considered for therapy in a wide range of diseases, including obesity, cancer, rheumatoid arthritis, asthma, and neurodegenerative diseases^48,50–54^. There are concerns with the toxic side-effects of Celastrol, which is an active area of development^48,55^. Both compounds reduced mutant mtDNA heteroplasmy and rescued mitochondrial fitness in the *uaDf5* animals, and Celastrol also improved several organismal fitness outcomes. The rescue observed upon Celastrol treatment was attenuated in the presence of RNAi against mitophagy genes, further demonstrating that the benefits of Celastrol treatment is at least partially conferred by modulation of mitophagy. As Celastrol modulates various pathways it may not be a direct activator of mitophagy^48^, which highlights the possibility that other biological pathways may play a role in improving the health of the *uaDf5* animals. Once identified, targeting these pathways may also open a path for therapeutic development in SLSMD disorders. Of note, finding highly variable effects based on sources of Hemin and Celastrol underscores the underlying variability that exists in quality of the drugs and natural products, even when compounds are obtained from the different production batches of the same supplier. Furthermore, rigorous, controlled clinical trials are needed to evaluate the safety and potential efficacy of Celastrol and Thiamine in human patients with SLSMD disorders.

Identification of Thiamine as an effective intervention in the *uaDf5* animals was surprising. It was identified in attempt to identify a positive experimental control compound for UPR^mt^ assay development in the *uaDf5* worms based on our research group’s having found it to be effective in other mitochondrial disease models^56,57^. Indeed, Thiamine (vitamin B1) is a commonly used and generally well-tolerated supplement that is used in some mitochondrial disorders such as those involving mitochondrial dehydrogenases, such as pyruvate dehydrogenase deficiency^58^. However, Thiamine has not previously been postulated to be a therapeutic modality for individuals with SLSMD disorders. Additional preclinical studies to understand the degree to which Thiamine rescues SLSMD animals’ complex organismal phenotypes need to be carried out along with a dissection of its underlying mechanism of action. Ultimately, given favorable benefit:risk balance of Thiamine, rigorous clinical trials may be considered to evaluate the potential therapeutic potential of high-dose Thiamine in individuals living with SLSMD disorders.

In summary, we have effectively utilized the *C. elegans* invertebrate animal model to study variable heteroplasmic levels in the SLSMD strain, *uaDf5,* both to gain deeper understanding of disease pathophysiology and to objectively screen for potential therapies. Screening a library of 62 mitophagy modulating compounds in *uaDf5* worms identified a positive control, Thiamine, as well as two candidate mitophagy modulating agents, Celastrol and Hemin, that significantly rescued mitochondrial unfolded protein (UPR^mt^) stress induction. Celastrol, a mitophagy activating anti-inflammatory and metabolic modifying natural product derived compound, rescued multiple fitness outcomes and reduced the SLSMD heteroplasmy level in *uaDf5* animals. This study highlights the utility of the *uaDf5* worm model to enable preclinical identification of therapeutic candidate leads for SLSMD-based heteroplasmic mtDNA diseases, along with the possible therapeutic potential of mitophagy modulators to improve health and reduce heteroplasmy in SLSMD diseases.

## ACKNOWLEDGEMENT

We thank Tali Gidalevitz, PhD, and Eiko Nakamaru-Ogiso, PhD, for helpful discussion and feedback. We would like to acknowledge Craig W. LaMunyon, PhD on helpful discussions on the *uaDf5* strain and providing additional aliquots of this worm strain. We would also like to thank the McGovern family and alongside the Satchel and CPEO/KSSS Impact Funds for raising funds for the United Mitochondrial Disease Foundation.

## AUTHOR CONTRIBUTION

SH and MJF conceived of the studies. SH trained and supervised RM, SM, AL. RM performed most experiments. SM performed the Gentle Touch Assay and the experiments (development, progeny count, and thrashing) that show diminishing defects over time. AL performed the thrashing experiments with Celastrol treatment. SH, RM, SM and AL performed the data analysis. SH and RM wrote the manuscript in consultation with MJF. All authors reviewed and approved the final manuscript.

