## Supplemental Figures for "Mitophagy modulation rescues single large-scale mitochondrial DNA deletion (SLSMD) disease symptoms in the *C. elegans uaDf5* animal model"

**SUPPLEMENTARY FIGURES**

**A. *uaDf5* strains and figures in which they are characterized**

| | wildtype genetics | designation | mutant genetics | designation | heteroplasmy |
|---|---|---|---|---|---|
| Fig S1 | N2 | N2 | *uaDf5* | | 0.4-5% |
| Fig 1 | *hsp-6p::GFP* | *hsp-6p::GFP* | *uaDf5; hsp-6p::GFP* | *uaDf5; hsp-6p::GFP #1* | <25% |
| Fig 1 | | | *uaDf5; hsp-6p::GFP* | *uaDf5; hsp-6p::GFP #2* | 25%-45% |
| Fig 2 | *myo-2p::mCherry; hsp-6p::GFP* | wildtype | *uaDf5; myo-2p::mCherry; hsp-6p::GFP* | Low Δ % | <25% |
| Fig 2 | | | *uaDf5; myo-2p::mCherry; hsp-6p::GFP* | Med Δ % | 25%-45% |
| Fig 2 | | | *uaDf5; myo-2p::mCherry; hsp-6p::GFP* | High Δ % | >45% |

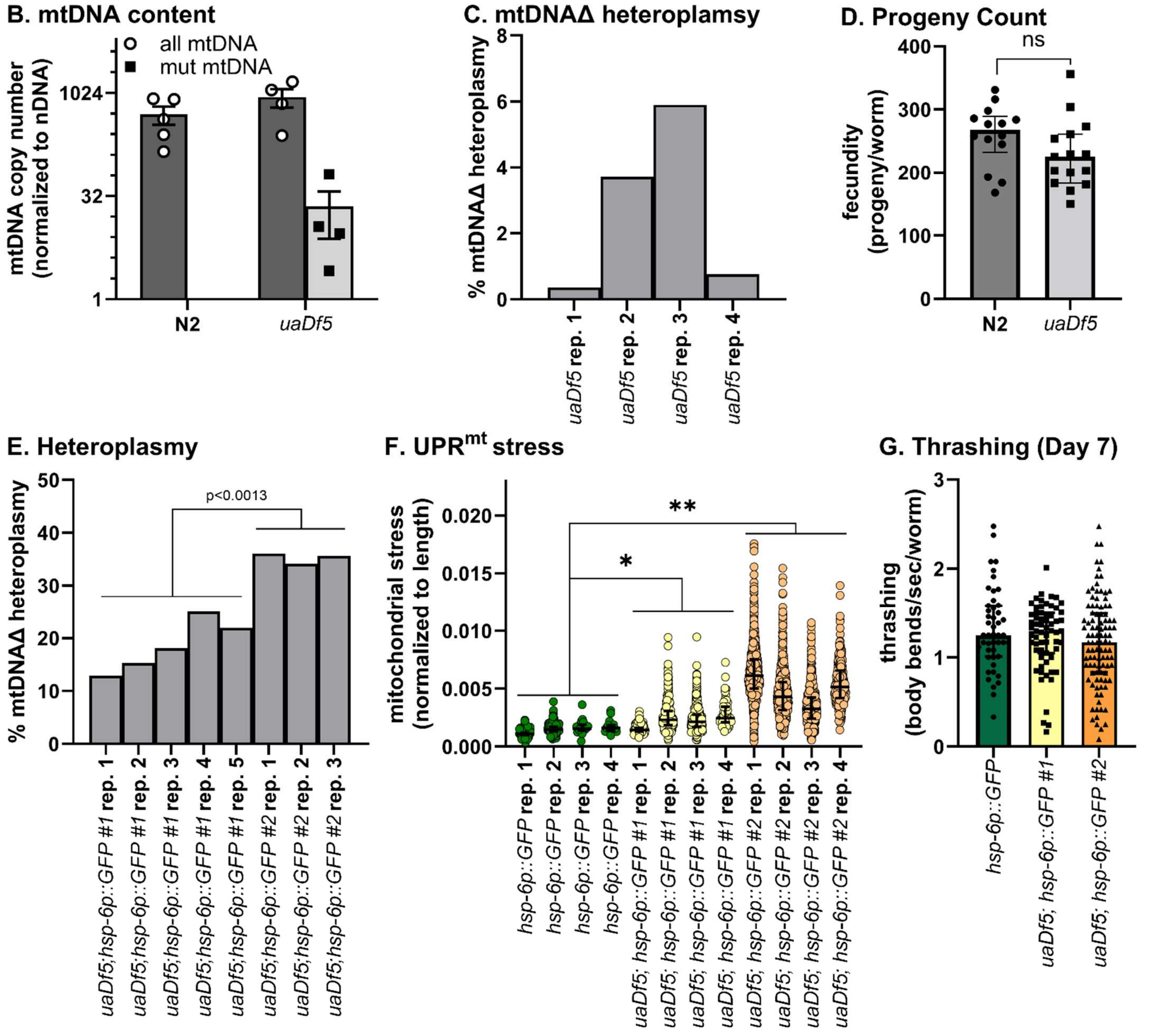


**Supplementary Figure 1: mtDNA assessment, development, and basal characterization of the *uaDf5* strains. (A)** Table of all strains in this study, including their genetics, designation, heteroplasmy levels, and the figure in which they are first mentioned. **(B)** Heteroplasmy assessment showed that the *uaDf5* animals had 2x total mtDNA (n.s.) as wildtype (N2) animals, and an average of 22 copies of mutant mtDNA per haploid nuclear genome (n.s.). **(C)** The heteroplasmy of each *uaDf5* sample in (B) was 0.37%, 3.7%, 5.6%, and 0.77%. **(D)** The *uaDf5* animals exhibited 16% decrease in fecundity (n.s.) compared to wildtype (N2) animals. **(E)** The heteroplasmy of each *uaDf;hsp-6p::GFP* #1 sample in Fig 1C were 13%, 15%, 18%, 25%, and 22%, and the heteroplasmy of each *uaDf;hsp-6p::GFP* #2 sample in Fig 1C were 36%, 34%, and 35%. **(F)** The individual replicate data of Fig 1C, assessing UPR$^{mt}$ using the COPAS Biosorter, showed that each wildtype sample (green)

exhibited stress at a median of 0.0011, 0.0015, 0.0016, 0.0016, each *uaDf;hsp-6p::GFP* #1 sample (yellow) exhibited stress at a median of 0.0015, 0.0023, o.0021, and 0.0025, and each *uaDf;hsp-6p::GFP* #2 sample (orange) exhibited stress at a median of 0.0061, 0.0043, 0.0033, 0.0052. **(G)** There was no neuromuscular defect observed when assessed via thrashing in the *uaDF5;hsp-6p::GFP #1* (yellow) and *uaDf5;hsp-6p::GFP #2* (orange) strains as compared to *hsp-6p::GFP* (green) animals at Day 7 of adulthood. For (B), each data point represents one biological replicate, 3-5 biological replicates were performed and the mean±SEM is graphed. For (C)(E), each bar represents one biological replicate. For (D)(G), the data was collected across 3 separate experiments and pooled where each data point represents the value per animal, and the median+interquartile range is graphed. For (F), each data point represents the value per worm and the median+interquartile range was graphed. Student's t-test analysis was performed to assess significance for (B)(D-G). (n.s.) applies to no statistical significance.

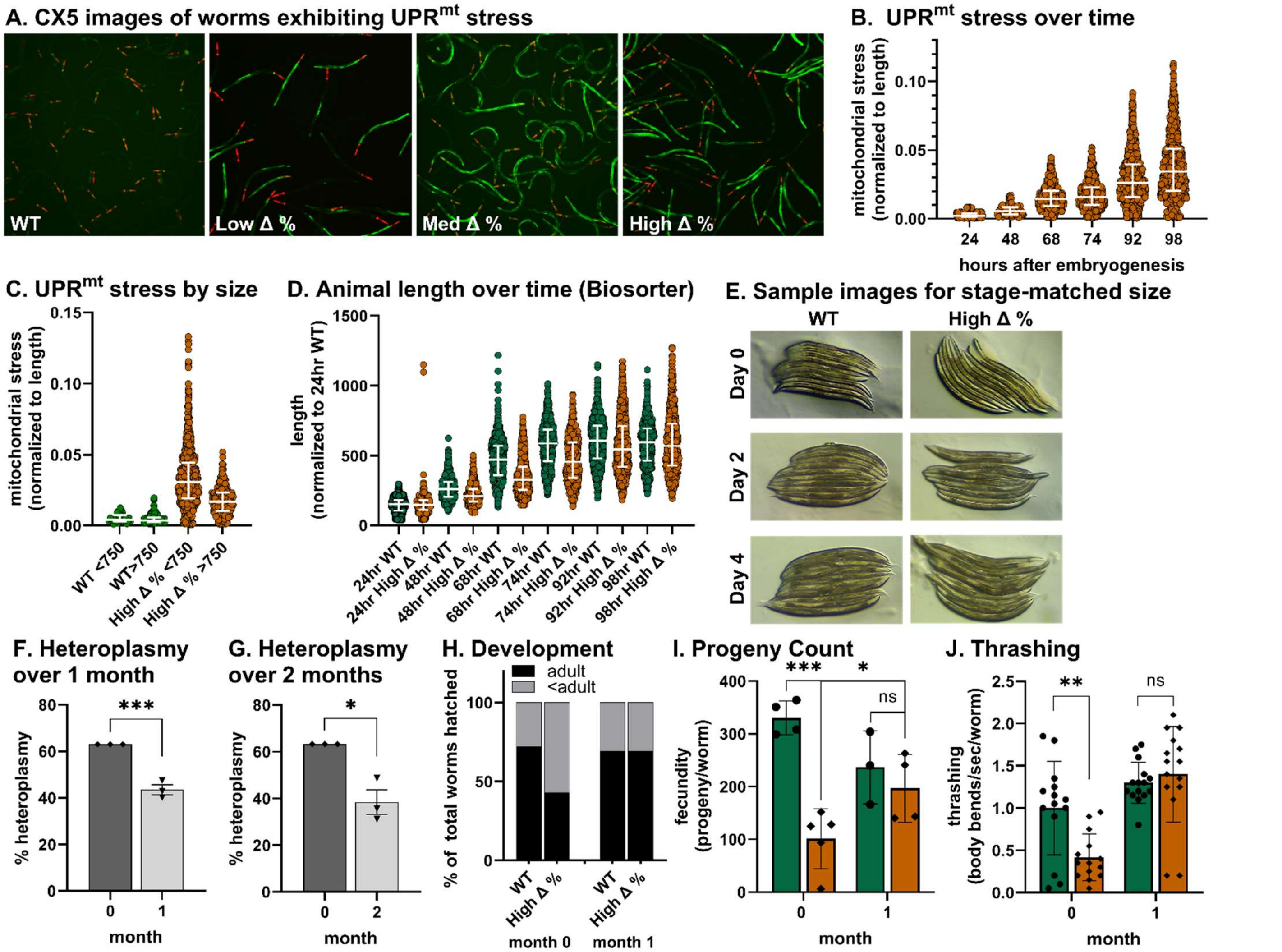


**Supplementary Figure 2: Characterization of *uaDf5* strains with *myo-2p::mCherry* and *hsp-6p::GFP* constructs. (A)** CX5 single well images of the WT, Low Δ %, Med Δ %, High Δ % animals, where the *myo-2p::mCherry* and *hsp-6p::GFP* reporters that confer red fluorescent heads for automated worm count and green fluorescence to measure $UPR^{mt}$. **(B)** $UPR^{mt}$ assessment at 24hr, 48hr, 68hr, 92hr, and 98hr post embryogenesis showed an increasing stress normalized to body length in the High Δ % animals. **(C)** Binning of the $UPR^{mt}$ assessment of animals at 98hr post embryogenesis by size of animals showed that the smaller (<750) animals exhibit a 1.8x higher $UPR^{mt}$ than the larger (>750) animals in the High Δ % strain (brown) and no difference in WT (green) animals. **(D)** Assessing size in WT (green) and High Δ % (brown) animals at 24hr, 48hr, 68hr, 92hr, and 98hr showed a 2%, 19%, 31%, 22%, 10% and 4.5% decrease in the High Δ % (brown) animals length compared to the WT (green) animals. **(E)** Images from replicate 1 of WT and High Δ % animals at L4, day 2 of adulthood and day 4 of adulthood used to assess size in stage matched animals. The % heteroplasmy in the High Δ % animals decreased by **(F)** 32% ($p<7.7x10^{-4}$) over one month and by **(G)** 40% ($p<0.0093$) over two months compared to the starting 63% heteroplasmy level. **(H)** Three days post embryogenesis, 43% of the animals reached adulthood in the High Δ % animals compared to 72% of animals reaching adulthood in WT animals in the same timeframe at month 0. The analysis of the same strains a month later showed that 69% of both mutant and WT strains reach adulthood 3 days post embryogenesis. **(I)** Fecundity assessment showed that High Δ % animals (orange) had an average of 101 progeny/worm, a 70% reduction ($p<1x10^{-4}$) compared to the 331 progeny/worm in WT animals (green). The analysis of the same strains a month later showed that mutant strain had an average of 197 progeny/worm, a 2x increase relative to the month 0 High Δ % animal progeny count. **(J)** The High Δ % (brown) strain exhibited 58% reduced thrashing rate at 3 days post embryogenesis ($p<0.002$) compared to WT (green) at month 0 and the same analysis a month later showed no difference between the WT (green) and mutant (brown) strains. For (B)-(D), each data represents values per worm from one biological replicate, and the median±interquartile range is graphed. For (F-G), each data represents values for a biological replicate, three biological replicate experiments were performed, and the mean+SEM is graphed. Single experiments were performed assessing 50 animals per condition in (H), 3-5 animals per condition in (I), and 14-16 animals per condition in (J). For (I-J), the mean±SD was graphed. Student's t-test analysis was performed to assess significance for (I-J).

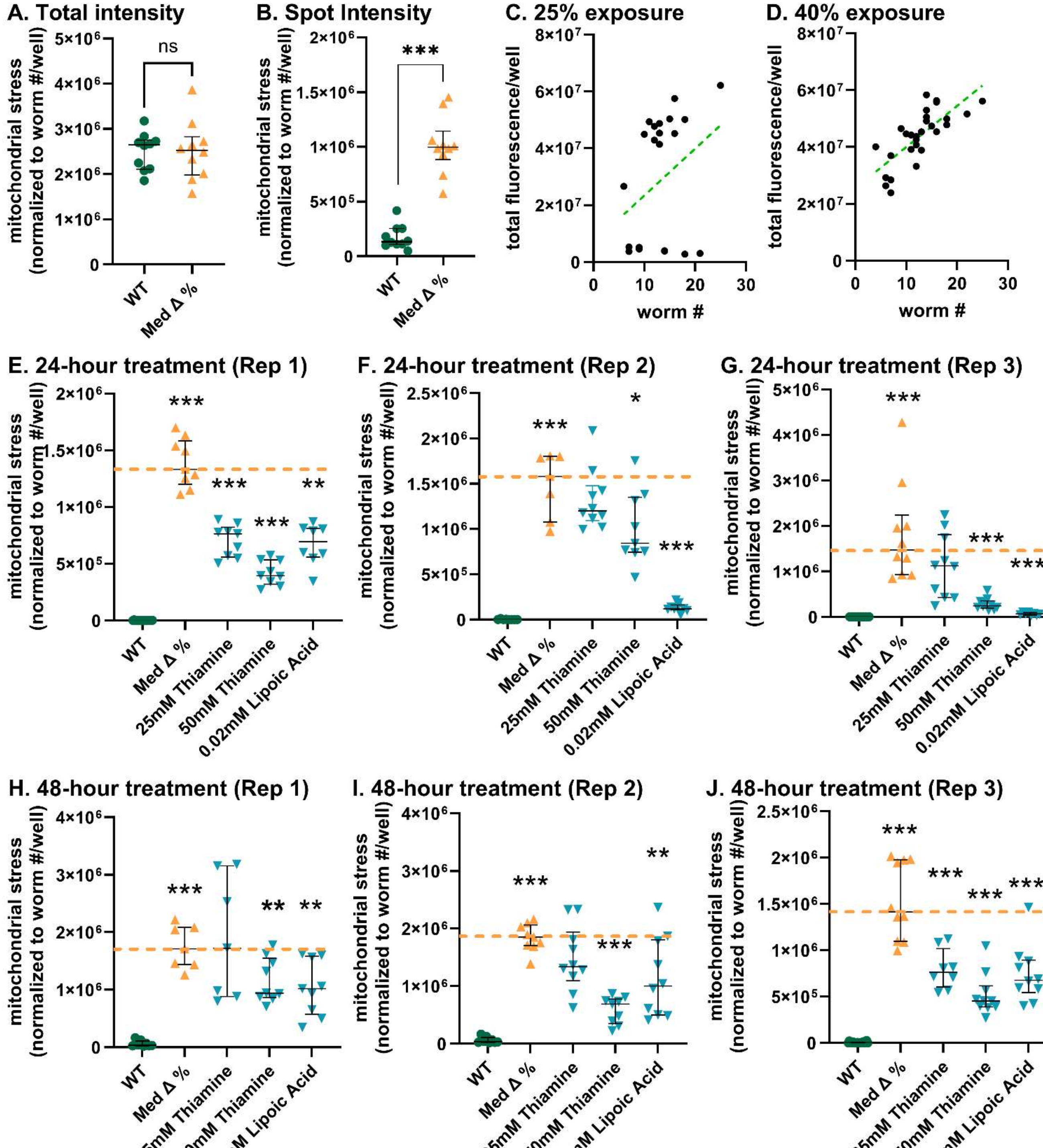


**Supplementary Figure 3: Developing screening protocol.** Two numerical outputs pertaining to the fluorescence per well outputted by the CX5 image analyzer; **(A)** total intensity output showed no difference in the expression of $UPR^{mt}$ between WT (green) and Med Δ % (orange) animals, and **(B)** spot intensity output showed that Med Δ % (orange) animals have a 5.8x times higher $UPR^{mt}$ ($p<2.3x10^{-8}$) compared to WT (green) animals. Threshold settings at **(C)** 25% exposure and **(D)** 40% exposure showed the 40% exposure recognizes each *cox-4p::cox-4::GFP* well to be fluorescent, whereas at 25% exposure, not all wells are captured to have fluorescent output. Biological replicate studies of 24-hour treatments show that 25mM Thiamine, 50mM Thiamine and 0.02mM Lipoic Acid reduced $UPR^{mt}$ by **(E)** 43% ($p<5.0x10^{-7}$), 70% ($p<1.6x10^{-9}$), and 48% ($p<1.9x10^{-6}$) in replicate 1, **(F)** 9.8% (n.s.), 37% ($p<0.025$), and 91% ($p<1.4x10^{-8}$) in replicate 2, and **(G)** 16% (n.s.), 82% ($p<3.1x10^{-4}$), and 95% ($p<3.3x10^{-4}$) in replicate 3. Biological replicate studies of 48-hour treatments showed that 25mM Thiamine, 50mM Thiamine and 0.02mM Lipoic Acid reduced $UPR^{mt}$ by **(H)** 0%, 45% ($p<0.0094$), and 40% ($p<0.0063$)

in replicate 1, **(I)** 22% (n.s.), 60% ($p<5.0x10^{-9}$), and 42% ($p<0.0095$) in replicate 2, and **(J)** 55% ($3.4x10^{-4}$), 73% ($p<2.4x10^{-6}$), and 60% ($p<1.4x10^{-4}$) in replicate 3. Each data point represents a per worm per well value (7-12 technical replicates) for (A)(B)(E-J) and the median$\pm$interquartile was graphed for the single biological replicate in each graph. Student's t-test analysis was performed to assess significance for (A, B, E-J). (n.s.) applies to no statistical significance.

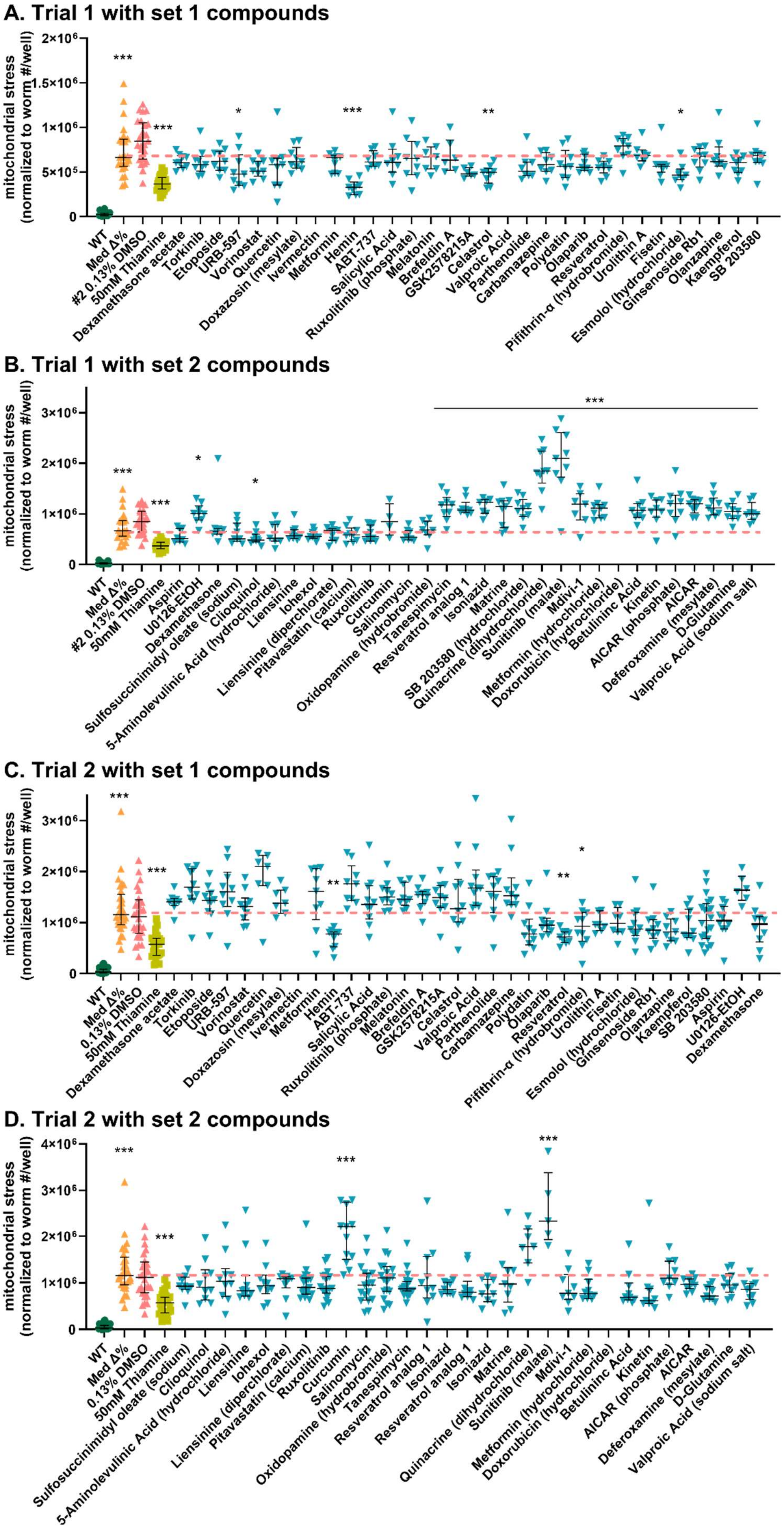
A. Trial 1 with set 1 compounds
mitochondrial stress (normalized to worm #/well)
2×10⁶
1.5×10⁶
1×10⁶
5×10⁵
0
***
***
*
***
**
*
WT
Med Δ%
#2 0.13% DMSO
50mM Thiamine
Dexamethasone acetate
Torkinib
Etoposide
URB-597
Vorinostat
Quercetin
Doxazosin (mesylate)
Ivermectin
Metformin
Hemin
ABT-737
Salicylic Acid
Ruxolitinib (phosphate)
Melatonin
Brefeldin A
GSK2578215A
Celastrol
Valproic Acid
Parthenolide
Carbamazepine
Polydatin
Olaparib
Resveratrol
Pifithrin-α (hydrobromide)
Urolithin A
Fisetin
Esmolol (hydrochloride)
Ginsenoside Rb1
Olanzapine
Kaempferol
SB 203580
B. Trial 1 with set 2 compounds
mitochondrial stress (normalized to worm #/well)
3×10⁶
2×10⁶
1×10⁶
0
***
***
***
*
*
WT
Med Δ%
#2 0.13% DMSO
50mM Thiamine
Aspirin
U0126-EtOH
Dexamethasone
Sulfosuccinimidyl oleate (sodium)
Clioquinol
5-Aminolevulinic Acid (hydrochloride)
Liensinine
Iohexol
Liensinine (diperchlorate)
Pitavastatin (calcium)
Ruxolitinib
Curcumin
Salinomycin
Oxidopamine (hydrobromide)
Tanespimycin
Resveratrol analog 1
Isoniazid
Matrine
SB 203580 (hydrochloride)
Quinacrine (dihydrochloride)
Sunitinib (malate)
Mdivi-1
Metformin (hydrochloride)
Doxorubicin (hydrochloride)
Betulininc Acid
Kinetin
AICAR (phosphate)
AICAR
Deferoxamine (mesylate)
D-Glutamine
Valproic Acid (sodium salt)
C. Trial 2 with set 1 compounds
mitochondrial stress (normalized to worm #/well)
3×10⁶
2×10⁶
1×10⁶
0
***
***
**
*
**
WT
Med Δ%
0.13% DMSO
50mM Thiamine
Dexamethasone acetate
Torkinib
Etoposide
URB-597
Vorinostat
Quercetin
Doxazosin (mesylate)
Ivermectin
Metformin
Hemin
ABT-737
Salicylic Acid
Ruxolitinib (phosphate)
Melatonin
Brefeldin A
GSK2578215A
Celastrol
Valproic Acid
Parthenolide
Carbamazepine
Polydatin
Olaparib
Resveratrol
Pifithrin-α (hydrobromide)
Urolithin A
Fisetin
Esmolol (hydrochloride)
Ginsenoside Rb1
Olanzapine
Kaempferol
SB 203580
Aspirin
U0126-EtOH
Dexamethasone
D. Trial 2 with set 2 compounds
mitochondrial stress (normalized to worm #/well)
4×10⁶
3×10⁶
2×10⁶
1×10⁶
0
***
***
***
***
WT
Med Δ%
0.13% DMSO
50mM Thiamine
Sulfosuccinimidyl oleate (sodium)
Clioquinol
5-Aminolevulinic Acid (hydrochloride)
Liensinine
Iohexol
Liensinine (diperchlorate)
Pitavastatin (calcium)
Ruxolitinib
Curcumin
Salinomycin
Oxidopamine (hydrobromide)
Tanespimycin
Resveratrol analog 1
Isoniazid
Resveratrol analog 1
Isoniazid
Matrine
Quinacrine (dihydrochloride)
Sunitinib (malate)
Mdivi-1
Metformin (hydrochloride)
Doxorubicin (hydrochloride)
Betulininc Acid
Kinetin
AICAR (phosphate)
AICAR
Deferoxamine (mesylate)
D-Glutamine
Valproic Acid (sodium salt)

**Supplementary Figure 4: Results of two replicate screens of the mitophagy modulator library.** The full screen of 62 mitophagy modulating compound library from medchemexpress was conducted across three experiments and repeated twice. **(A)(B)** In the first replicate of the screen, the Med Δ % animals (orange) exhibited 28x more $UPR^{mt}$ ($p<6.7x10^{-8}$) relative to WT animals (green), show no statistical difference to DMSO treated animals (pink), and 50mM Thiamine treatment (light green) reduced $UPR^{mt}$ by 45% ($p<6.7x10^{-7}$). **(A)** 25µM URB-597, Hemin, Celastrol, and Esmolol (hydrochloride) treatment reduced $UPR^{mt}$ by 28% (p<0.046), 50% ($p<1.9x10^{-4}$), 25% (p<0.012), and 30% (p<0.020) respectively. **(B)** 25µM Clioquinol treatment reduced $UPR^{mt}$ by 28% (p<0.038). 25µM U0126-EtOH and compounds from Tanespimycin to Valproic Acid treatment increased $UPR^{mt}$ by 1.5x (p<0.012) and 1.5-2.8x ($p<5.0x10^{-10}$-p<0.001). **(C)(D)** In the second replicate of the screen, the Med Δ % animals (orange) exhibited 35x more $UPR^{mt}$ ($p<4.4x10^{-12}$) relative to WT animals (green), showed no difference to DMSO treated animals (pink), and 50mM Thiamine treatment (light green) reduced $UPR^{mt}$ by 50% ($p<2.0x10^{-8}$). **(C)** 25µM Hemin, Resveratrol, and Pifithrin-α (hydrobromide) treatment reduced $UPR^{mt}$ by 33% (p<0.0017), 38% (p<0.0036), and 20% (p<0.038) respectively. **(D)** 25µM Curcumin, and Sunitinib (malate) treatment increased $UPR^{mt}$ by 1.9x ($p<1.5x10^{-4}$) and 2.0x ($p<4.4x10^{-5}$). Each data point represents 6-11 technical replicate wells per experiment. The data for the control conditions (WT, Med Δ %, 0.13% DMSO and 50mM Thiamine) were combined across the three separate experiments caried out per library screen. The median±interquartile range is graphed, and student's t-test analysis was performed to assess significance.

| | Replicate | | | Replicate | |
|---|---|---|---|---|---|
| **Drug** | **1** | **2** | **Drug** | **1** | **2** |
| Dexamethasone acetate | black | black | Aspirin | black | black |
| Torkinib | black | black | U0126-EtOH | green | black |
| Etoposide | black | black | Dexamethasone | black | black |
| URB-597 | green | black | Sulfosuccinimidyl oleate (sodium) | black | black |
| Vorinostat | black | black | Clioquinol | green | black |
| Quercetin | black | black | 5-Aminolevulinic Acid (hydrochloride) | black | black |
| Doxazosin (mesylate) | black | black | Liensinine | black | black |
| Ivermectin | black | black | Iohexol | black | black |
| Metformin | black | black | Liensinine (diperchlorate) | black | black |
| Hemin | green | green | Pitavastatin (calcium) | black | black |
| ABT-737 | black | black | Ruxolitinib | black | black |
| Salicylic Acid | black | black | Curcumin | black | red |
| Ruxolitinib (phosphate) | black | black | Salinomycin | black | black |
| Melatonin | black | black | Oxidopamine (hydrobromide) | black | black |
| Brefeldin A | black | black | Tanespimycin | red | black |
| GSK2578215A | black | black | Resveratrol analog 1 | red | black |
| Celastrol | green | black | Isoniazid | red | black |
| Valproic Acid | black | black | Matrine | red | black |
| Parthenolide | black | black | SB 203580 (hydrochloride) | red | black |
| Carbamazepine | black | black | Quinacrine (dihydrochloride) | red | red |
| Polydatin | black | black | Sunitinib (malate) | red | black |
| Olaparib | black | green | Mdivi-1 | red | black |
| Resveratrol | black | green | Metformin (hydrochloride) | red | black |
| Pifithrin-α (hydrobromide) | black | black | Doxorubicin (hydrochloride) | red | black |
| Urolithin A | black | black | Betulininc Acid | red | black |
| Fisetin | black | black | Kinetin | red | black |
| Esmolol (hydrochloride) | green | black | AICAR (phosphate) | red | black |
| Ginsenoside Rb1 | black | black | AICAR | red | black |
| Olanzapine | black | black | Deferoxamine (mesylate) | red | black |
| Kaempferol | black | black | D-Glutamine | red | black |
| SB 203580 | black | black | Valproic Acid (sodium salt) | red | black |

**Supplementary Figure 5: Table displaying the outcome of the duplicate mitophagy modulator library screen.** The table summarizing data presented in **Supp. Fig 4**, where the compounds in orange are soluble in DMSO, the compounds in blue are soluble in water, green boxes indicate statistically significant decrease (rescue) of the $UPR^{mt}$, red boxes indicate statistically significant increase of the $UPR^{mt}$, and black boxes indicate no statistical difference between treated and untreated animals. Student's t-test analysis was performed on data displayed in **Supp. Fig 4** to assess significance reported in this table.

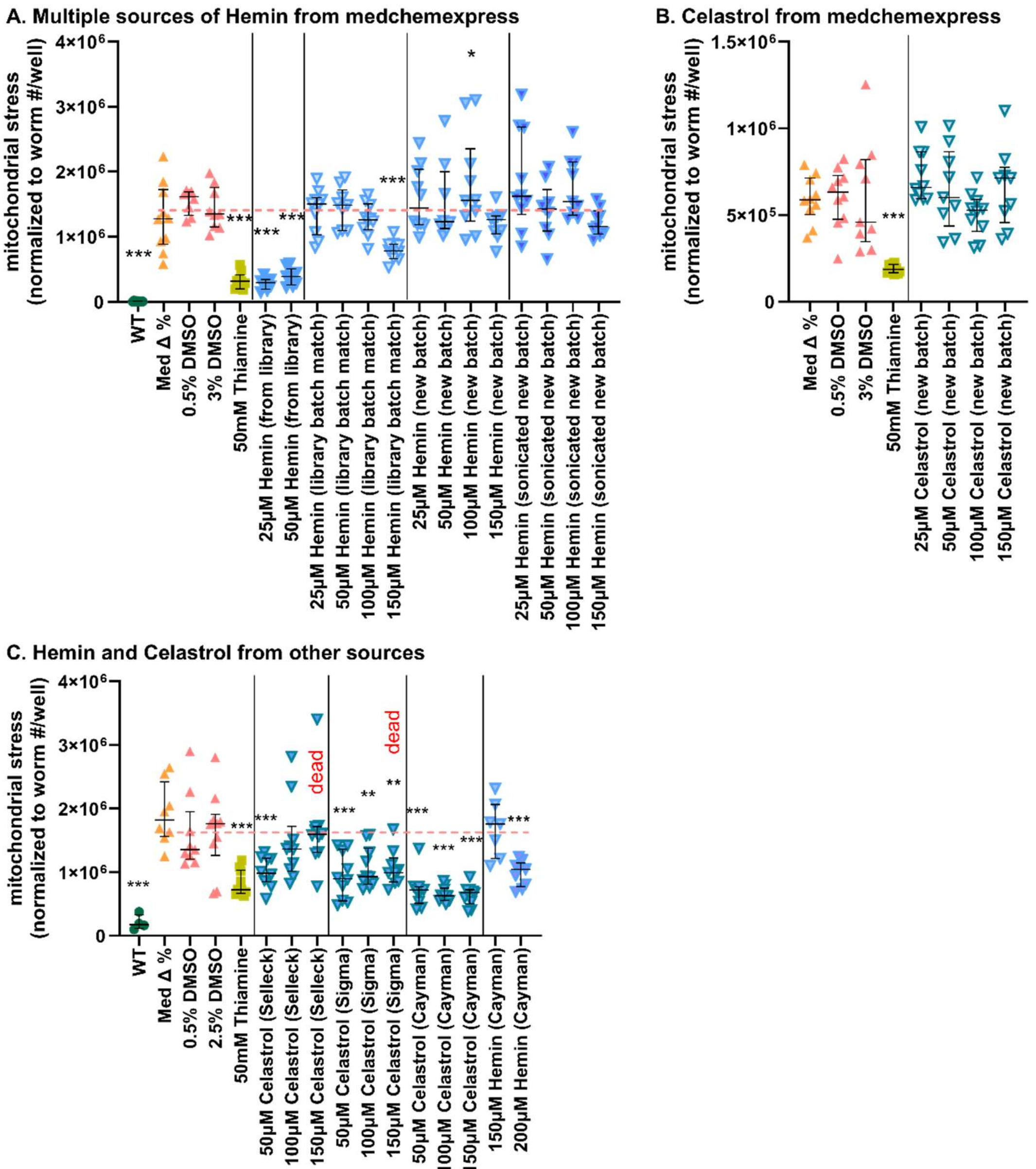


**Supplementary Figure 6: Testing different sources of Hemin and Celastrol. (A)** The Med Δ % animals (orange) exhibited 177x more $UPR^{mt}$ ($p<5.7\times10^{-11}$) relative to WT animals (green), and showed no statistical difference to 0.5% and 3% DMSO treated animals (pink), and the 50mM Thiamine treatment (light green) reduced $UPR^{mt}$ by 77% ($p<1.0\times10^{-9}$). 25μM and 50μM Hemin from the library rescued $UPR^{mt}$ by 80% ($p<1.7\times10^{-9}$) and 73% ($p<4.0\times10^{-9}$) respectively. 150μM Hemin from medchemexpress with the same batch number as the Hemin provided in the library reduced $UPR^{mt}$ by 45% ($p<1.7\times10^{-5}$). 100μM Hemin from medchemexpress with a different batch number from the Hemin provided in the library increased $UPR^{mt}$ by 1.3x ($p<0.049$). Sonicated Hemin from medchemexpress with a different batch number from the Hemin provided in the library has no significant effect on $UPR^{mt}$. **(B)** 50mM Thiamine treatment (light green) reduced $UPR^{mt}$ by 68% ($p<2.6\times10^{-8}$) and Celastrol from medchemexpress with a different batch number from the Celastrol provided in the library had no effect on $UPR^{mt}$. **(C)** The Med Δ % animals (orange) exhibited 9x more $UPR^{mt}$ ($p<5.0\times10^{-5}$) relative to WT animals (green) and showed no statistical difference to 0.5% and 2.5% DMSO treated animals (pink), and the 50mM Thiamine treatment (light green) reduced $UPR^{mt}$ by 52% ($p<3.4\times10^{-5}$). 50μM Celastrol from Selleck reduced $UPR^{mt}$ by 41% ($p<6.4\times10^{-4}$). 50, 100, and 150μM Celastrol from Sigma decreased $UPR^{mt}$ by 45% ($p<3.6\times10^{-4}$), 37% ($p<0.0021$), and 58% ($p<0.0027$) respectively. 50, 100, and 150μM Celastrol from Cayman reduced $UPR^{mt}$ by 58% ($p<7.4\times10^{-6}$), 62% ($p<1.6\times10^{-6}$), and 63% ($p<1.4\times10^{-6}$). 200μM Hemin from Cayman reduced $UPR^{mt}$ by 43% ($p<7.2\times10^{-4}$). 150μM Celastrol (from Selleck and Sigma) treatment resulted in dead animals on the day of assessment. Each data point represents 6-11 technical replicate wells per experiment and the median$\pm$interquartile range is graphed. Student's t-test analysis was performed to assess significance.
